# Social Touch Suppresses Aggression via Thalamic Mechanisms

**DOI:** 10.1101/2025.08.23.670446

**Authors:** Tamás Láng, Botond B. Drahos, Fanni Dóra, Dávid Keller, Ingrid Csordás, Vivien Szendi, Gina Puska, Valery Grinevich, Árpád Dobolyi

## Abstract

Understanding the neural circuitry underlying aggression is critical for both scientific insight and clinical intervention. Here, we identify the posterior intralaminar thalamic nucleus (PIL) as a key node in an anti-aggressive circuit activated by social touch. Using a rodent model, we demonstrate that deprivation of direct physical contact during social isolation leads to heightened aggression. Chemogenetic and optogenetic manipulations reveal that PIL neurons activated by social touch inhibit aggression via excitatory projections to the medial preoptic area (MPOA) of the hypothalamus. This PIL-to-MPOA pathway is suppressed by input from the ventromedial hypothalamic nucleus (VMH). Our findings establish a novel thalamic-hypothalamic circuit that mediates social touch-induced suppression of aggression, offering potential targets for therapeutic intervention in conditions marked by pathological aggression.

## Introduction

Understanding the neural mechanisms of aggressive behavior is essential for both scientific insight and clinical application. Aggression, while a natural and adaptive behavior that evolved to protect resources, defend territory, and establish social hierarchies, can become maladaptive when expressed in excessive, inappropriate, or uncontrolled forms (*1*). Considerable progress has been made in identifying subcortical regions that drive aggressive behavior. The ventromedial hypothalamic nucleus (VMH) has been firmly established as a pivotal node facilitating intermale aggression in rodents (*2–4*). Stimulation of this region can elicit attack behaviors even in the absence of social provocation, while its inhibition can abolish naturally occurring aggression. However, the VMH does not act in isolation, and the complete architecture of brain regions that regulate aggression, especially those that suppress it, remains poorly understood. One intriguing line of evidence comes from studies on sensory modulation of social behavior. A recent report demonstrated that the cortical amygdaloid nucleus in mice functions as an inhibitory center of aggression, relying heavily on olfactory inputs to gate hostile responses (*5*). However, in humans, gentle tactile contact such as a hug, caress, or stroking has been observed to reduce aggressive tendencies, often accompanied by increased prosocial communication. This phenomenon has inspired therapeutic interventions, including massage therapy, for aggression reduction in children and adolescents, as well as in elderly individuals experiencing dementia-related agitation (*6*). Despite the clinical application of touch-based therapies (*7*), the neural pathways through which social touch inhibits aggression remain ill defined. Recent work in rodents has identified the medial preoptic area (MPOA) of the hypothalamus as a recipient of gentle tactile input, with this activity modulating the quality and intensity of social interactions (*8*). The MPOA is recognized for its role in reproductive, parental, and affiliative behaviors. Its capacity to integrate multimodal sensory signals makes it a candidate for mediating the anti-aggressive effects of social touch (*9*). Indeed, the MPOA was shown to influence male aggressive behaviors (*2*) via its projections to the VMH (*10*).

Previous studies have identified the posterior intralaminar nucleus (PIL), which is located medially within the lateral thalamus, as a critical sensory relay that transmits salient somatosensory and multimodal information to the forebrain (*11*). The PIL has previously been implicated in the control of maternal behaviors (*12–14*) as well as auditory fear response (*15*). Furthermore, PIL neurons are responsive to social cues (*16*). As previously demonstrated, PIL neurons projecting directly to the MPOA and that PIL neurons exhibit c-Fos activation in response to social touch. Furthermore, activation of the PIL-to-MPOA pathway promotes social grooming in female rats, suggesting a role in reinforcing affiliative contact (*17*). In light of these findings, we hypothesized that the same PIL-to-MPOA circuitry engaged by affiliative tactile input might also modulate aggressive behavior. Specifically, we predicted that activation of this pathway would inhibit intermale aggression by shifting behavioral priorities toward prosocial or affiliative actions. To test this hypothesis, a series of experiments were conducted on rats and mice to determine the functional contribution of the PIL-to-MPOA pathway to aggression control, using behavioral, activity-mapping, opto-, and chemogenetic approaches, as well as gene expression analysis.

### Aggression and social isolation

Male Wistar rats housed in groups rarely show aggression toward an unfamiliar male intruder. However, extensive prior research has demonstrated that prolonged social isolation can markedly increase aggressive responses in rodents (*18–21*). Consistent with this literature, we found that two weeks of complete isolation in a standard rat cage was sufficient to induce pronounced aggressive behavior toward a male intruder in the home cage. The isolated rats displayed frequent attack postures, vigorous chasing, and biting directed at the intruder within minutes of its introduction. Next, we sought to determine whether aggression develops in the absence of somatosensory social contact or it can be prevented by maintaining other channels of social information. To this end, we developed a partial social deprivation model in which two rats were housed in the same cage but were separated by a fixed metal grating (Fig. 1A). This barrier prevented direct tactile contact between the animals while permitting continuous exchange of visual, auditory, and olfactory cues. The aggressive behavior of these animals measured in a resident intruder test was compared to that of a group of animals kept in complete social isolation through single housing. A third, control group was pair-housed in an identical cage without a barrier, allowing unrestricted tactile and close-proximity interactions. These rats frequently engaged in mutual grooming, huddling, and sleeping in physical contact. Behavioral assessments revealed striking differences between the groups over the course of the one-month experimental period in the resident intruder test performed at the beginning of the experiment (day 0) when all animals used to be housed three per cage, two weeks later, and four weeks later (Fig. 1A). Control rats with unrestricted contact maintained low aggression levels, comparable to baseline, and predominantly exhibited affiliative and exploratory behaviors toward an introduced conspecific. In contrast, the barrier-housed rats deprived of social touch, despite retaining visual, auditory, and olfactory communication, developed aggressive responses that were statistically indistinguishable from those of fully isolated rats (Fig. 1B). These animals readily attacked an unfamiliar intruder, displaying similar attack latency, bite frequency, and aggressive bout duration as the solitary group. These findings demonstrate that tactile contact is indispensable for maintaining low aggression levels in male Wistar rats. The absence of social touch, even in the presence of other sensory modalities, is sufficient to trigger an aggression-prone phenotype. This suggests tactile interactions play a unique and non-redundant role in regulating aggression-related neural circuits. Alternative sensory channels appear insufficient to compensate for the behavioral buffering effects of somatosensory social contact.

**Fig. 1.**
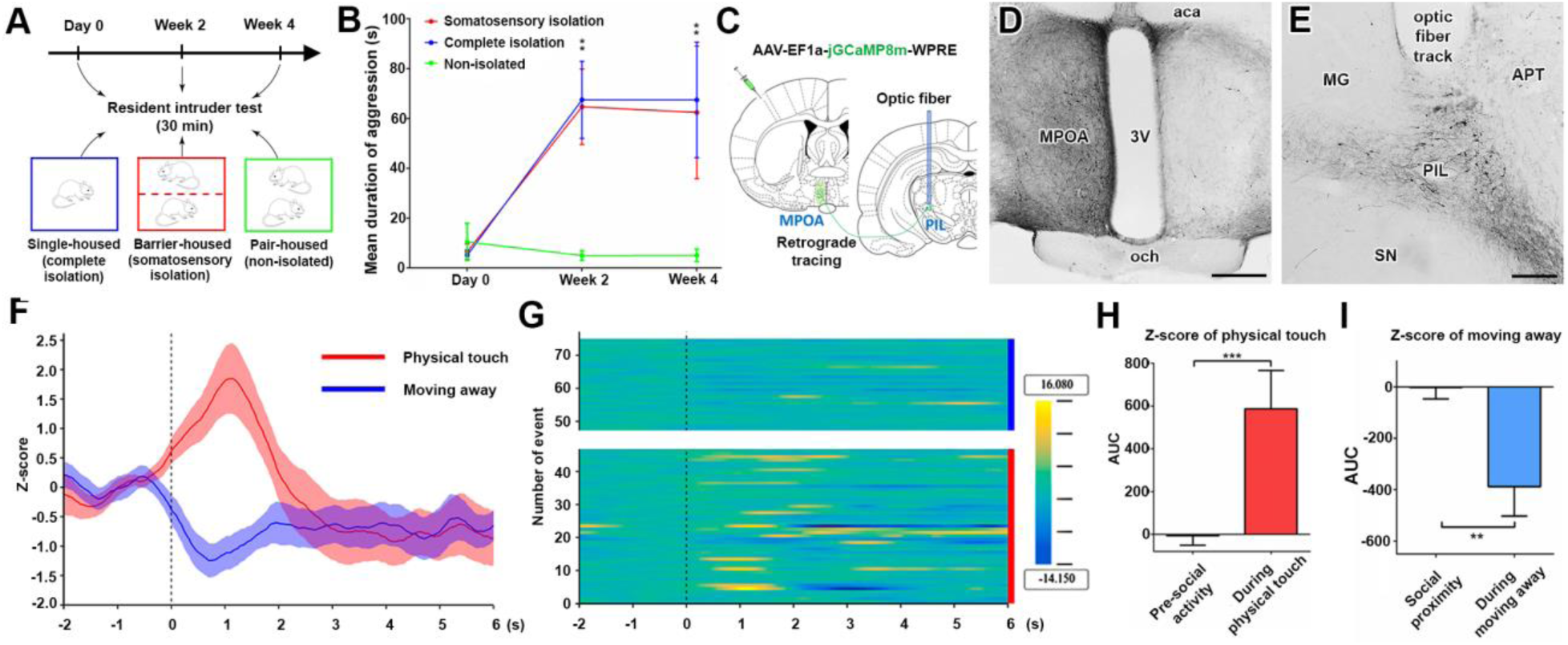
Behavioral and brain activation consequences of social touch. **(A)** The schematic of the experimental design is presented, illustrating the allocation of the animals into three groups and the designated time points for evaluation in the resident intruder test. (**B**) Consequences of prolonged complete social isolation and somatosensory social isolation on the subjects. Pair-housed rats (green line) demonstrated no increase in aggression throughout the experiment. In contrast, fully isolated animals (blue) and animals deprived of social touch (red) exhibited increased aggression to a similar degree. (**C**) A schematic figure illustrates the surgical protocol for fiber photometry. The MPOA was targeted with a retrogradely spreading virus expressing green fluorescent protein (GCaMP) (AAV-EF1a-jGCaMP8m-WPRE), and optic fibers were implanted above the PIL. (**D**) The histological image of the injection site in the MPOA is presented herein. The scale bar is 500 µm. (**E**) The image illustrates the retrogradely labeled cell bodies and the precise location of the optic cannula above the PIL. The scale bar is 250 µm. (**F**) The mean and standard error of the mean (SEM) of calcium activity Z-scores of PIL neurons projecting to the MPOA during physical contact (visualized in red) and when moving away (visualized in blue) are presented. The 0-time stamp on the X-axis delineates the onset of the behavioral elements. Furthermore, a phyasic increase in calcium activity was observed that persisted for a duration of two seconds. This increase commenced at the onset of the behavior, defined as the physical contact or movement away from the subject. Z-scores of deltaF/F values are displayed on the y-axis. (**G**) The Z-scores for each event are displayed in a heat map. The heat map illustrates a total of 75 events, of which 47 (0–46) are designated as physical touch (bottom, marked with a vertical red line at the right) and 28 (47–75) are marked as moving away (top, marked with a vertical blue line at the right). The scale of calcium activity is displayed on the right side of the image. (**H**) Statistical analysis of social touch: The two-second Z-score AUC was then compared to the two-second Z-score AUC from the previous two seconds, which corresponded to the moment of social touch. (**I**) A statistical analysis of the phenomenon of moving away was conducted. The 2-second Z-score area under the curve (AUC) was compared to the AUC of the Z-scores 2 seconds of social proximity prior to moving away.

### Posterior thalamic neurons projecting to the MPOA are activated by social touch

The MPOA has been demonstrated to modulate social homeostasis through distinct neuronal populations that respond to social isolation and social touch, respectively (*8*). Retrograde neuronal tract tracing of MPOA neurons, coupled with c-Fos activation studies induced by social interaction, suggests that the MPOA receives information about social touch sensation from the PIL (*17*). The activity of the MPOA-projecting PIL neurons was measured with fiber photometry. An adeno-associated virus (AAV) with retrograde propagation encoding the calcium sensor protein GCaMP8 was injected into the MPOA while an optic fiber guide was implanted immediately above the PIL (Fig. 1C). The virus infected several fiber terminals in the MPOA (Fig. 1D), and expressed GCaMP in neuronal cell bodies in the PIL (Fig. 1E), which projected to the MPOA. Following the surgical procedure and a recovery period in isolation, the subjects were placed in an arena with a conspecific for the purpose of free interaction. Subsequently, the activity of the PIL neurons was measured. In the context of these conditions, no instances of aggression were observed among the animals. An increase in fluorescence (z-score of deltaF/F) occurred for approximately two seconds when the animals touched each other, and the z-score decreased as they moved away (Fig. 1F,G). No other behavioral element demonstrated a significant correlation with PIL neuron activity. For instance, during the periods of pre-social activity and social proximity, the area under the z-score curve remained unchanged (Fig. 1H,I). These results indicate that prosocial physical touch activates PIL neurons that project to the MPOA.

### Neurons in the PIL inhibit aggressive behavior

To investigate the function of PIL neurons in aggression, we employed a chemogenetic approach by injecting adeno-associated viruses (AAVs) that expressed designer receptors exclusively activated by designer drugs (DREADDs) into the nucleus (Fig. 2A). The stimulation of PIL neurons led to a reduction in the aggressive behavior exhibited by male rats in response to a male intruder placed within their home cage, as compared to both the previous and the subsequent control test day. Concurrently, the duration of positive valence contact (grooming, side-to-side contact, friendly mounting) exhibited an increase, while other social behaviors not involving contact, such as body and anogenital sniffing, remained unaltered (Fig. 2B1). Given the possibility of chemogenetic stimulation to elicit supraphysiological effects, in another experiment, the physiological activity of the PIL neurons was chemogenetically inhibited. This protocol resulted in a substantial escalation in aggression (Fig. 2C1). The application of the receptor-activating chemical agent clozapine-N-oxide (CNO) to rats expressing a control chemogenetic receptor resulted in no observable effect (Fig. 2D1). The results of chemogenetic stimulation and inhibition suggest a physiological role of PIL neurons in reducing the motivation for aggression. A subsequent analysis of the types of aggressive behavior altered by stimulation and inhibition revealed that dominant mounting and fighting were all affected with biting also showing a tendency of change suggesting a general aggression-reducing effect of the PIL output neurons (Fig. 2B-D). To circumvent the possibility of nonspecific effects resulting from altered locomotion or anxiety levels induced by manipulating PIL neurons, an elevated plus maze test was conducted in conjunction with the social interaction experiment. The parameters measured in this test, including the number of entries into the arms (Fig. 2B4-D4) and the time spent in the arms (Fig. S1A-C), remained constant in response to manipulation of PIL neuronal activity, implying that the observed changes in aggression are not a consequence of alterations in locomotor activity or anxiety levels.

**Fig. 2.**
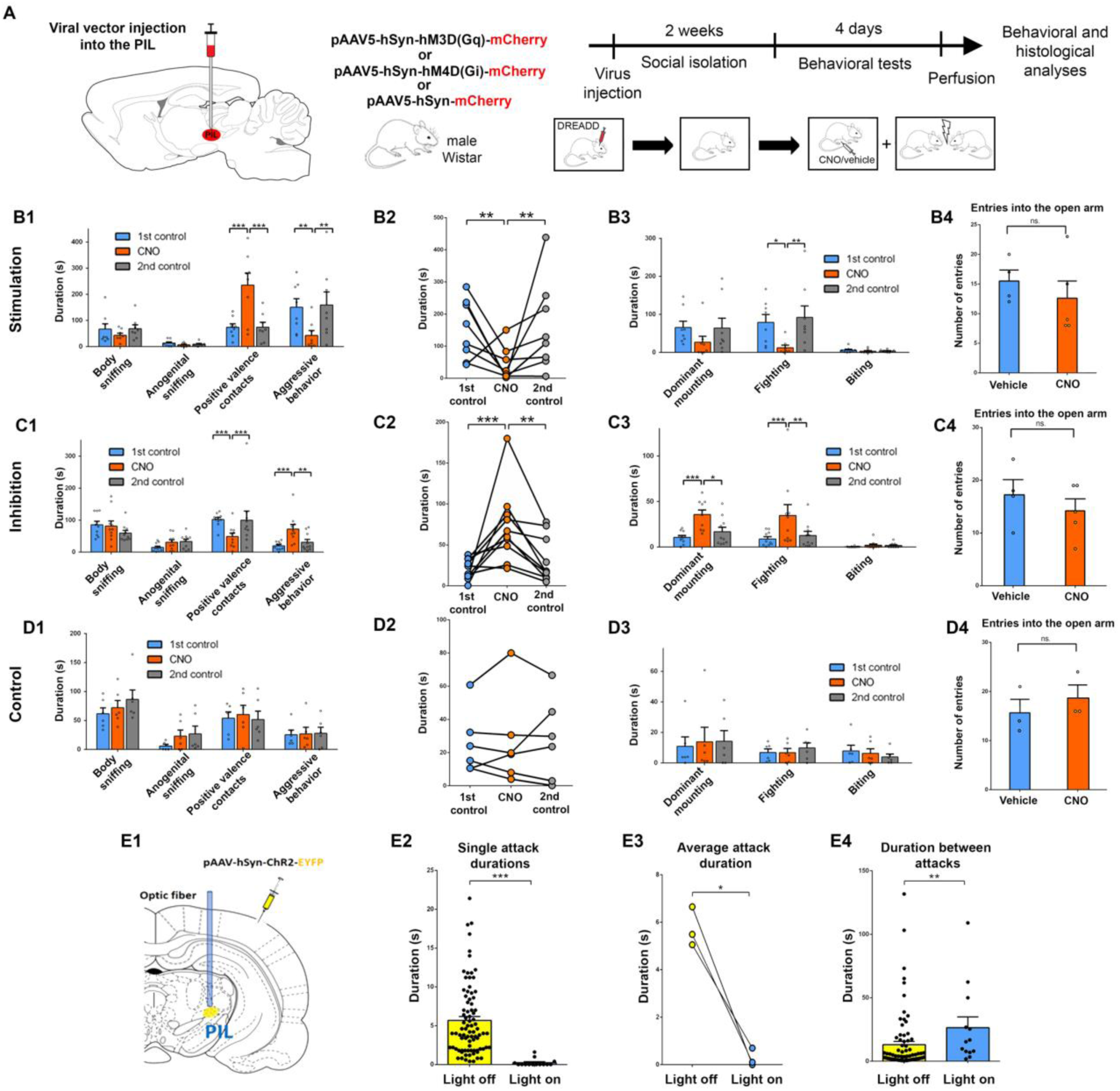
The effect of the stimulation and inhibition of PIL neurons on aggression. (**A**) A schematic representation illustrates the viruses injected into the PIL, in conjunction with the protocol for testing aggression in response to chemogenetic manipulation. (**B1**) The chemogenetic stimulation of PIL neurons resulted in a substantial reduction in the duration of aggressive behavior. (**B2**) The panel demonstrates a significant reduction in aggressive behaviors exhibited by the animals during the chemogenetic stimulation of PIL neurons. (**B3**) Within the aggression category, mounting and fighting decreased significantly in response to the chemogenetic stimulation of PIL neurons. (**B4**) The chemogenetic stimulation of PIL neurons does not affect locomotor activity in the elevated plus maze paradigm. (**C1**) The application of chemogenetic inhibition to PIL neurons resulted in a decline in positive valence contact duration and an escalation in aggression. Statistical significance was identified in both behavioral elements. (**C2**) The panel demonstrates the heightened aggressive behavior exhibited by the animals during the chemogenetic inhibition of PIL neurons. (**C3**) Within the aggression category, mounting and fighting increased significantly in response to the chemogenetic inhibition of PIL neurons. (**C4**) The chemogenetic inhibition of PIL neurons does not result in alterations to locomotor activity. (**D1**) The administration of CNO did not elicit any observable effects on the behavior of animals that had been injected with the control virus. (**D2**) The aggressive behavior exhibited by the animals was not influenced by CNO in subjects that had been injected with the control virus into the PIL. (**D3**) CNO does not modify any behavioral elements of aggression. (**D4**) The administration of CNO did not result in alterations in the locomotor activity of the animals in the elevated plus maze paradigm. (**E1**) The schematic design of the viral injection into the PIL and the placement of the optic fiber in CD1 male mice. (**E2**) Photostimulation of PIL neurons during aggressive behavior was demonstrated to result in a dramatic reduction in the duration of individual attacks. (**E3**) Optogenetic stimulation of PIL neurons during aggressive behavior was demonstrated to markedly reduce the mean duration of attacks. (**E4**) A long-term consequence of optogenetic stimulation during an attack is an augmentation of the duration prior to the occurrence of the subsequent attack.

Given the inability of chemogenetics to rapidly modulate the activity of affected neurons, we also employed optogenetics to ascertain whether stimulating PIL neurons could effectively halt aggression. The injection of an AAV encoding a stimulatory opsin into the PIL was followed by the implantation of an optic fiber above the targeted area (Fig. 2E1). In this study, wild-type CD1 mice were used because, unlike Wistar rats they are sufficiently aggressive in an arena outside of the home cage. Turning on the light for 0.5 seconds at the start of an attack reduced its duration from 5.73±0.48 seconds to 0.27±0.22 seconds (Fig. 2E2,3). This stimulation had a long-lasting effect, as subsequent attacks were delayed; they occurred 26.43±8.53 seconds after the previous attack, as opposed to the 13.08±2.59 seconds inter-attack interval when the light was off (Fig. 2E4). The results suggest that heightened activity of the PIL neurons can promptly halt aggressive behavior. Furthermore, the sustained neuronal activation exerts a persistent influence on the timing of subsequent attacks.

### Socially-activated PIL neurons are sufficient for the aggression-reducing effect

Chemogenetic receptors were next specifically expressed in the PIL neurons, which were activated in response to social interaction with another male rat. To this end, the PIL was injected with a viral cocktail (Fig. 3A), designated vGATE, which has been previously utilized to successfully express DREADDs exclusively in neurons that are c-Fos-activated during a time window permitted by antibiotic administration (6). This study demonstrates that social interaction elicits c-Fos expression in specific PIL neurons in male rats (Fig. 3B,C). Consequently, the expression of stimulatory and inhibitory DREADDs was observed in these neurons (Fig. 3D). During social interaction, the animals predominantly exhibited positive-valence contact. A significant proportion of time was dedicated to body sniffing, and instances of aggressive behavior were also documented (Fig. 3E). Following an additional 10 days of social isolation, the animals underwent the male intruder test, during which they exhibited aggressive behavior (Fig. 3D), which was inhibited by stimulation of the socially-tagged PIL neurons (Fig. 3F-H). Conversely, a significant escalation in aggressive behavior was documented in response to the inhibition of socially-tagged PIL neurons (Fig. 3I-K). This finding suggests a causal relationship in which the activity of these neurons is sufficient to inhibit aggression. The projections of the socially-tagged neurons were established through the tracing of the fluorescent protein mCherry fused with the DREADD. The major projection sites of these PIL neurons from posterior to anterior include the periaqueductal grey, dorsomedial hypothalamic nucleus, medial amygdaloid nucleus, paraventricular hypothalamic nucleus (PVN), MPOA, the ventral subdivision of the lateral septal nucleus, and the infralimbic cortex (Fig. S2).

**Fig. 3.**
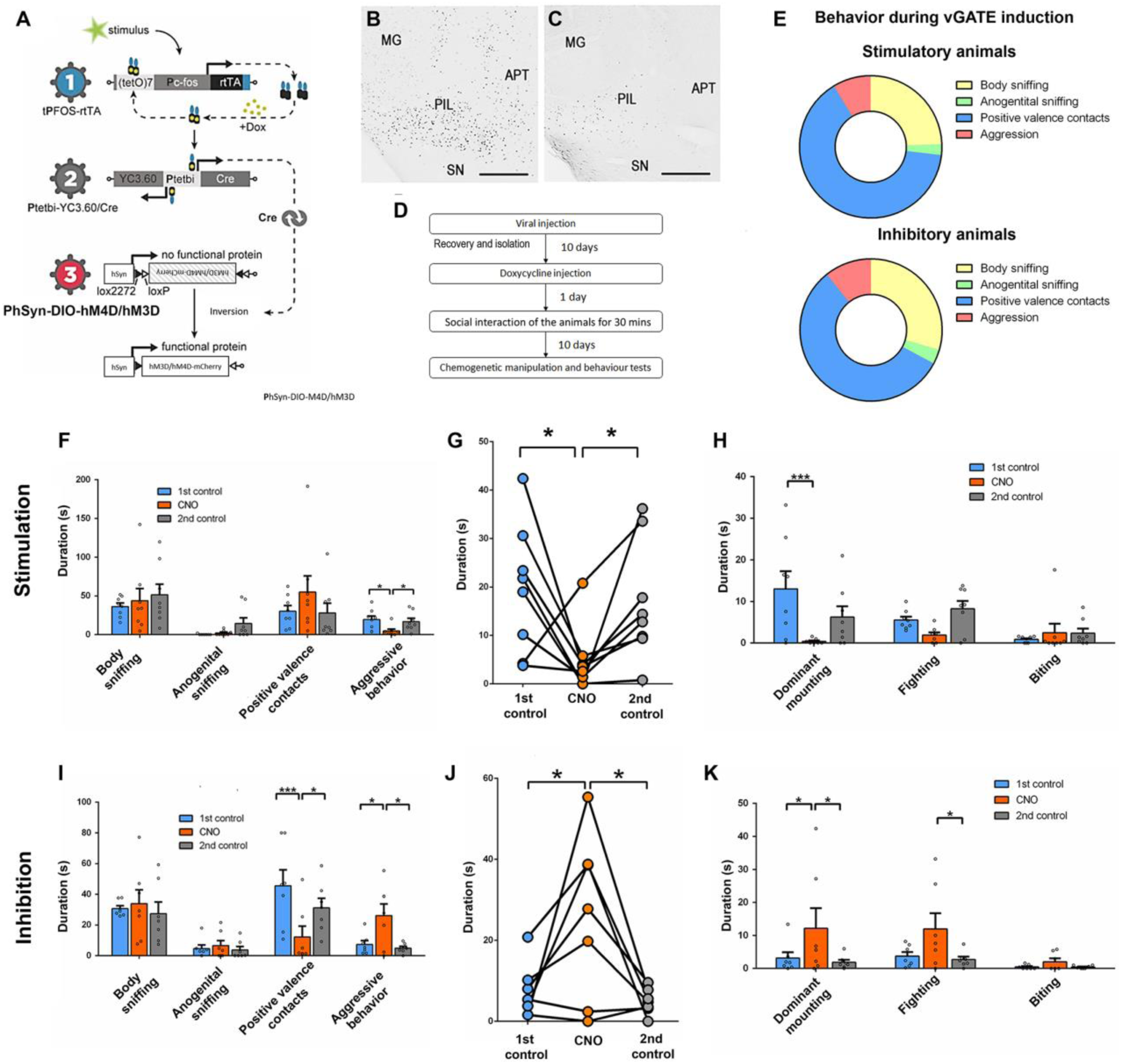
Social activity tagged PIL neurons are sufficient and necessary for the inhibition of aggression. (**A**) A schematic representation of the vGATE (Virus-delivered Genetic Activity-induced Tagging of Cell Ensembles) system is presented, revealing the mechanism by which a combination of three viruses i) (rAAV-(tetO)7-CAG-GC-hCFC-IRES-mCherry, ii) rAAV-Ptet-Cre/YFP, and iii) rAAV-Phsyn-DIO-hM3D(Gq)-mCherry or rAAV-Phsyn-DIO-hM4D(Gi)-mCherry, injected into the PIL results in the expression of DREADD in c-Fos-activated neurons (*50*). (**B**) Social interaction between unfamiliar male rats, in which aggression is present, was demonstrated to induce c-Fos expression in the PIL. (**C**) In the control group, consisting of socially isolated animals, c-Fos was not expressed in PIL neurons. Scale bar: 500 µm for panels B and C. (**D**) The protocol that was employed in the study was designed to tag socially activated PIL neurons. The viral mixture was administered into the PIL. During the 10-day recovery period, the animals were subjected to social isolation. The activation of the vGATE system was then initiated through the administration of doxycycline to the animals, followed by the execution of the resident-intruder test the subsequent day. Subsequent to a further 10-day period of anticipation, during which the DREADDs were expressed in adequate quantities, behavioral tests were conducted on the animals employing chemogenetic manipulation. (**E**) The exhibition of social behavior by the animals that expressed stimulatory and inhibitory DREADDs during the induction of the vGATE system subsequent to the doxycycline injection was observed. (**F**) The chemogenetic stimulation of socially-tagged PIL neurons led to a significant decrease in the time spent engaging in aggressive behavior. (**G**) Individual data concerning the animals’ aggressive behaviors during chemogenetic stimulation of socially-tagged PIL neurons. (**H**) In the context of aggression, mounting and fighting exhibited a substantial decrease in response to the chemogenetic stimulation of socially-tagged PIL neurons. **(I**) The application of chemogenetic inhibition to socially-tagged PIL neurons was demonstrated to result in a decline in positive valence contact duration and an escalation in aggression. Statistical significance was identified in both behavioral elements. (**J**) Individual data concerning the animals’ aggressive behaviors during chemogenetic inhibition of social-tagged PIL neurons. (**K**) In the context of aggression, mounting and fighting increased significantly in response to the chemogenetic inhibition of social-tagged PIL neurons.

### Projections of PIL neurons to the MPOA convey an aggression-reducing effect

Chemical excitation of PIL neurons by CNO administration in rats previously injected with a virus expressing stimulatory DREADD exhibited increased c-Fos expression in the infected neurons as compared to the vehicle injected control animals providing evidence that the stimulatory action of the DREADD is functioning (Fig. S3A,C). Furthermore, heightened c-Fos expression was detected in the MPOA of these rats (Fig. S3B,D). However, this was not observed in other brain regions (Fig. 4A), indicating that PIL neurons predominantly stimulate neurons in the MPOA among their target areas.

**Fig. 4.**
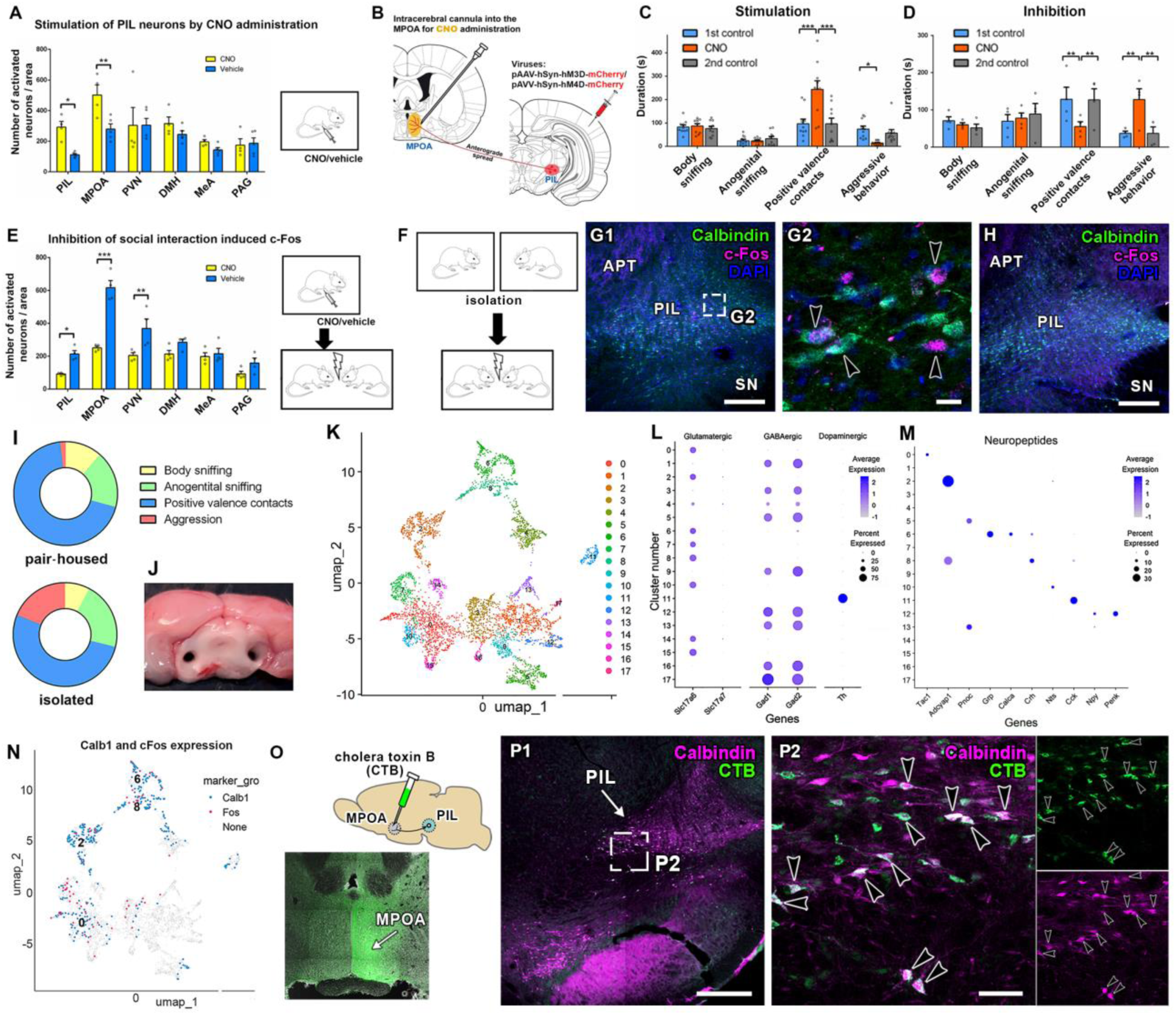
Cell types within the PIL, which may convey social touch information to the MPOA. (**A**) A substantial number of c-Fos-expressing neurons was identified in the PIL following chemogenetic stimulation of PIL neurons. Elevated c-Fos expression in the medial preoptic area (MPOA) was observed in response to chemogenetic stimulation of PIL neurons in the absence of social interaction. (**B**) The schematic figure illustrates the experimental design for manipulating the PIL-to-MPOA pathway. (**C**) The impact of chemogenetic stimulation of the PIL-to-MPOA pathway on behavioral responses to a male intruder. (**D**) The impact of chemogenetic inhibition of the PIL-to-MPOA pathway on behavioral responses to a male intruder. (**E**) A quantitative analysis was conducted on c-Fos-immunoreactive neurons in the PIL and its target brain areas. Subsequent to chemogenetic inhibition of PIL neurons, a marked diminution in activity was observed within the PIL and MPOA during social interactions. (**F**) The experimental design for the c-Fos and calbindin immunohistochemical analysis. The rats were separated for 22 hours and then reunited for 90 minutes before perfusion. (**G1**) The low-magnification image of a brain section containing the PIL demonstrates a high number of activated neurons in the area where calbindin neurons are located. Scale bar: 100 µm. Abbreviations: APT – anterior pretectal nucleus, PIL - posterior intralaminar thalamic nucleus, SN - substantia nigra. (**G2**) A higher magnification of the double-labeled c-Fos and calbindin-positive neurons (arrowheads) in the PIL is presented. Scale bar: 20 µm. (**H**) Rats that were separated without reunion were used as controls and did not show significant c-Fos positivity in the PIL. Scale bar: 100 µm. **(I)** Behavioral differences between pair-housed and socially isolated rats. Donut charts show the proportion of time spent on body sniffing, anogenital sniffing, positive valence contacts, and aggressive behaviors. (**J**) Representative image of the dissected posterior intralaminar thalamic nucleus (PIL) used for single-nucleus RNA sequencing. (**K**) UMAP plot of the re-clustered neuronal population from pooled PIL samples, showing 18 transcriptionally distinct subclusters. (**L-M**) Dot plots represent the average expression and proportion of cells expressing selected neurotransmitter-related and neuropeptide genes across neuronal clusters. (**N**) UMAP plot highlighting clusters containing c-Fos-positive and Calb1-positive cells. (**N**) A quantitative analysis was conducted on c-Fos-immunoreactive neurons in the PIL and its target brain areas. Subsequent to chemogenetic inhibition of PIL neurons, a marked diminution in activity was observed within the PIL and MPOA during social interactions. (**O**) The experimental design and the injection site of the retrograde tract-tracing of the MPOA by using the retrogradely transporting cholera toxin B subunit (CTB) in rat. (**P1**) The distribution of retrogradely labeled CTB positive neurons (green) overlap with the distribution of Calbindin-ir neurons (purple) in the medial part of the PIL. Scale bar: 100 µm. (**P2**) A higher magnification photomicrograph showing double labeled CTB- and calbindin-positive neurons (white) in the medial part of the PIL. Some double-labeled neurons are pointed to by black arrowheads. The right panels display the green and purple channels separately for better visibility of their labeling. Scale bar: 50 µm.

The role of the PIL-to-MPOA pathway in aggression was subsequently investigated with chemogenetics. Stimulatory and inhibitory DREADDs were expressed in the PIL via AAV injections, and a low concentration of CNO, and vehicle injection as control, was applied locally into the MPOA via previously implanted bilateral guide cannulae (Fig. 4B). Local stimulation of the MPOA terminals of PIL neurons led to reduced aggression (Fig. 4C), while their inhibition resulted in elevated aggression (Fig. 4D), suggesting the role of the PIL-to-MPOA pathway in controlling aggressive behavior. A general rather than a specific effect on aggression can be inferred from the finding that the duration of dominant mounting changed significantly, but the duration of fighting and biting also showed strong tendencies in the same direction (Fig. S4).

The PIL, and the brain regions to which the socially-activated, vGATE tagged PIL neurons project, exhibited elevated levels of c-Fos activation in response to a male intruder (Fig. S5). The chemogenetic inhibition of the PIL led to a reduction in social interaction-induced c-Fos activation in PIL neurons (Fig. S3E,G), thereby demonstrating the effectiveness of the inhibitory DREADD expressed in the PIL neurons. Additionally, the social interaction-induced c-Fos was also reduced in the MPOA (Fig. S3F,H) but not in most other brain regions (Fig. 5E). This finding indicates that the input from the PIL substantially contributes to the social interaction-induced activation of MPOA neurons.

**Fig. 5.**
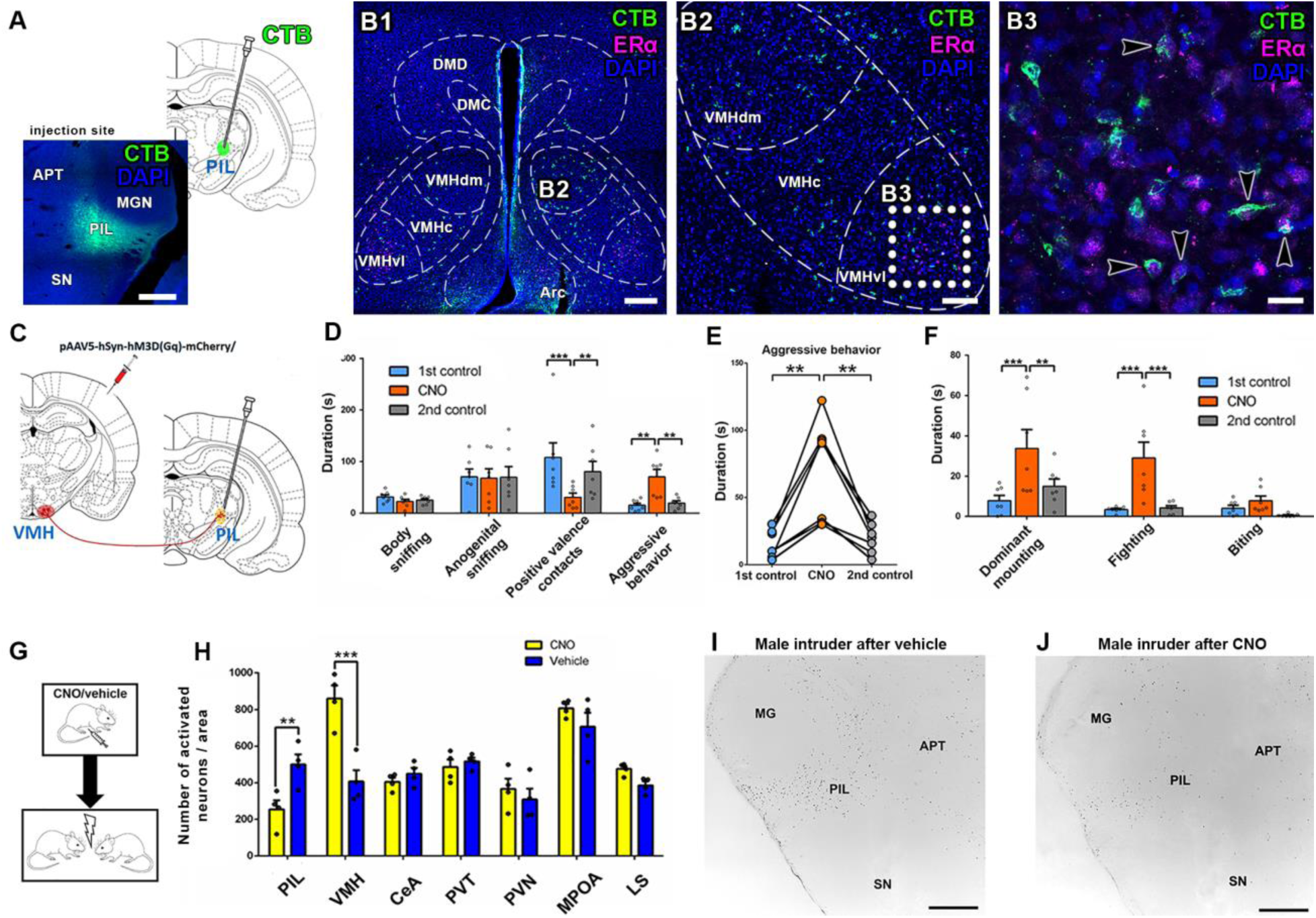
VMH input to the PIL eliminates the aggression-reducing action. (**A**) Injection site of cholera toxin beta (CTB) in the PIL. . Scale bar: 500 µm (**B1**) Retrogradely labelled (CTB-positive, green) neurons in the tuberal region of the hypothalamus. Scale bar: 250 µm. Abbreviations: Arc – arcuate nucleus, DMC – dorsomedial hypothalamic nucleus, compact part, DMD – dorsomedial hypothalamic nucleus, dorsal part, VMHc – ventromedial hypothalamic nucleus, central part, VMHdm – ventromedial hypothalamic nucleus, ventrolateral part, VMHdm – ventromedial hypothalamic nucleus, dorsomedial part. (**B2**) The location of retrogradely labeled neurons in the ipsilateral VMH is distributed in all subdivisions. Scale bar: 100 µm. (**B3**) The high magnification confocal photomicrograph demonstrates that only a portion of the retrogradely labeled neurons is estrogen receptor alpha (ERα)-positive (arrowheads) in the ventrolateral subdivision of the VMH, the area where ERα neurons are abundant. Scale bar: 20 µm. (**C**) The schematic figure illustrates the experimental design for manipulating the VMH-to-PIL pathway. The VMH was targeted with stimulatory DREADD-expressing viral vectors (pAAV-hSyn-hM3D(G_q_)-mCherry), and intracerebral cannulas were implanted above the PIL for selective chemogenetic stimulation of the terminals of the pathway. (**D**) Chemogenetic stimulation of the VMH-to-PIL pathways resulted in heightened aggression and diminished positive valence contacts. (**E**) The individual data regarding aggression in response to chemogenetic stimulation of the VMH-to-PIL pathway. (**F**) Among aggressive behavioral elements, dominant mounting and fighting increased significantly with biting showing a similar tendency. (**G**) Schematics of the c-Fos study. The animals were administered either CNO or a vehicle control, and subsequently exposed to a male intruder. (**H**) A quantitative analysis of c-Fos expression in VMH and its target areas during VMH stimulation was conducted. VMH demonstrated considerably elevated levels of activity, while PIL exhibited a conspicuously diminished level of activity. (**I**) Male intruder-induced c-Fos activity appears in PIL neurons following vehicle injection. (**J**) In response to stimulation of the input from the VMH by local CNO injection, PIL neurons exhibited reduced activation under the same conditions. Abbreviations: APT – anterior pretectal nucleus, MG – medial geniculate body, PIL - posterior intralaminar thalamic nucleus, SN - substantia nigra.

### PIL cell types mediating the aggression-reducing effects

#### Calbindin neurons in the PIL project to the MPOA and are activated by aggression

Neurons containing the calcium-binding protein calbindin were abundant in the PIL but rare in adjacent regions. When double-labeled with c-Fos in male rats exposed to a male home cage intruder, a large portion of the c-Fos-activated neurons were found to contain calbindin (Fig. 4F-H) indicating the significance of PIL calbindin neurons in mediating social touch information from the PIL to the MPOA.

#### PIL region single-nucleus RNA sequencing and quality metrics

To address the number of different neuronal cell types in the PIL and whether calbindin neurons constitute a single cell type, and which cell types are c-Fos-activated, single-nucleus RNA sequencing (snRNA-seq) of the PIL was performed in three groups: 1) pair-housed controls, which were separated only for 24 h before dissection to reduce basal socially-induced c-Fos activity (group Cont), 2) animals pair-housed until one day before the dissection, then separated and reunited with their previous cage mate one hour before sacrifice to evoke affiliative interaction-induced c-Fos expression (group Affi) (Fig. 4I), 3) animals isolated for a period of two weeks and subsequently exposed to an unfamiliar home-cage intruder for one hour prior to euthanasia, with the objective of studying c-Fos expression following aggressive behavior (group Aggr) (Fig. 4I). The PIL was subsequently dissected bilaterally using micropunch technique (*22*) from the 12 male rats (Fig. 4J). Samples from the four animals within the same experimental group were combined to provide sufficient material for snRNA-seq of the 3 groups. We retained 26,918 high-quality nuclei (Cont: 7,794; Affi: 7,983; Aggr: 11,141). Median gene counts per nucleus ranged from 1,155 to 1,556, and median UMI counts from 1,822 to 2,880, indicating robust transcript capture (Table S1). Violin plots visualizing the distribution of gene counts (nFeature_RNA), unique molecular identifier (UMI) counts (nCount_RNA), and mitochondrial gene percentages (percent.mt) across samples are shown in Fig. S6A. Uniform Manifold Approximation and Projection (UMAP) embedding colored by sample origin showed no significant batch effects (Fig. S6B). Following dimensionality reduction and unsupervised clustering, we identified 19 transcriptionally distinct clusters comprising both neuronal and non-neuronal cell types identified by canonical marker genes (Table S2 and Fig. S6C). The final UMAP embedding colored by cell identity highlighted this cellular diversity within the PIL (Fig. S6D).

#### Neuronal subclusters and their characterization

To identify neuron-specific transcriptional programs associated with distinct behavioral states, we focused our analysis on the neuronal subset of the PIL dataset. Unsupervised clustering of all high-quality nuclei initially yielded 23 neuronal clusters. To eliminate sampling bias from the variation of the position of dissections, we applied a stringent inclusion criterion: only 13 clusters in which all three experimental groups contributed at least 10% of total neurons were retained. Neurons in these clusters were then re-integrated and re-clustered to capture finer transcriptomic distinctions within the neuronal population. This re-clustering step, performed at higher resolution, produced 18 transcriptionally distinct neuronal subclusters, as visualized in a UMAP embedding (Fig. 4K) with distinct genes characterizing them (Table S3 and Fig. S7A).

The identification of the neurotransmitters of these 18 neuronal PIL clusters revealed that 8 of them were glutamatergic, 8 of them GABAergic, 1 dopaminergic, and 1 expressed markers of both glutamatergic and GABAergic neurons (Fig. 4L). The presence of additional neuropeptide transmitters was observed in six glutamatergic (clusters 0,2,6,7,8,10), three GABAergic (clusters 5,12,13), and the dopaminergic cluster (cluster 11) (Fig. 4M, Table S4). Dopaminergic cells are not present in the PIL but rather likely derive from the ventrally adjacent substantia nigra. Therefore, they were not considered in subsequent analyses.

Neuropeptides are characterized by their highly restricted expression patterns and relatively high levels of expression. This characteristic renders them particularly well-suited for the study of the spatial distribution of a specific neuronal cluster in which they are expressed. Consequently, a comprehensive search was conducted in the Allen Brain Atlas to ascertain the distribution of the mRNA of these neuropeptides within the PIL established by *in situ* hybridization histochemistry (*23*). The peptidergic clusters exhibited distinct distribution patterns within the PIL, with only a subset of them demonstrating significant spatial overlap with each other (Fig. S8).

#### Behavioral activation of c-Fos^+^/Calb1^+^ neurons in the PIL

To identify which of the PIL neuronal clusters may be involved in transmitting signal on social touch, we analyzed calbindin and c-Fos expression based on the above-described histological data that the majority of social interaction-induced c-Fos cells contain calbindin (Calb1) (Fig. 4F-H). It was first established that the Calb1-expressing neurons predominantly belong to 4 clusters (0,2,6,8) containing the highest numbers of Calb1-positive neurons. Combined, they contain 305 out of 397 Calb1-positive neurons (76.8%). Interestingly, all 4 clusters were glutamatergic in nature. In the UMAP plot of the integrated dataset, the distribution of c-Fos and Calb1 expression was similar (Fig. 4N, Fig. S7B), therefore, we analyzed c-Fos expression in the 3 experimental groups (Cont, Affi and Aggr) in these 4 clusters (Table S5). Based on Chi square test, the ratio of c-Fos-positive neurons in both the Affi and Aggr groups (15 and 31 out of 328 and 985, respectively) was higher (p <0.05) than in the control group (7 out of 705) while Affi and Aggr groups were not statistically different from each other. These data suggest that calbindin-expressing, glutamatergic neurons in these 4 clusters play a major role in conveying information on social interaction.

To further identify which cells activate MPOA neurons in social contexts, we compared the distribution of the four clusters obtained by neuropeptide content (Fig. S8) with the distribution pattern of neurons projecting from the PIL to the MPOA. The latter was determined by injecting the retrograde tracer cholera toxin B subunit into the MPOA. The retrogradely labeled neurons within the PIL were located medially (Fig. 4O,P). In this region, 83.5±6.4% (mean±SD) of them co-localized with calbindin while calbindin neurons in the lateral part of the PIL lacked retrograde labeling. The medial location of double-labeled neurons corresponds to neurons belonging to clusters 0 and 2 (Fig. S8A1,2). In contrast, neurons belonging to clusters 6 and 8 are located laterally within the PIL, suggesting that these neurons do not project to the MPOA. These findings provide substantial evidence in support of the hypothesis that neurons in clusters 0 and 2 convey information about social touch to the MPOA.

#### Communication profile of MPOA-projecting socially activated PIL neurons reveal high integrative capacity

To understand how MPOA-projecting socially activated PIL neurons integrate within the nucleus, we investigated the intercellular communication profiles of PIL neuronal subclusters using cell-cell communication analysis (*24*). To quantify global signaling potential, we calculated total incoming and outgoing interaction strengths per cluster (Fig. S9A). Clusters 0 and 2 received similar incoming signals, suggesting a comparable inter-nuclear connectivity and potentially analogous functions. Their incoming signals are strong and balanced, implying an integrative role in processing socially relevant inputs. Both are characterized mainly by pattern 3 type input (Fig. S9B), which consists primarily of semaphorin 3 (SEMA3), and to a lesser extent bone morphogenetic protein (BMP), and neuregulin (NRG) signaling pathway inputs. Since we only examined connections between neurons, the Sema3, BMP, and NRG input signals likely originate from neurons in other PIL clusters belonging to the output heatmap patterns 1 and 4 (Fig. S9C). As for Sema3, the inputs originate from clusters 10 and 15. A similarly selective structure emerged in the outgoing communication space (Fig. S9C). Clusters 2 and 6 showed strong alignment with neurotrophin (NT), and insulin-like growth factor (IGF)-centered outgoing programs, indicating potential roles in providing trophic support to downstream targets. In the Sankey diagram of latent pattern decomposition, Calb1⁺ clusters were among the most prominent contributors to a dominant communication program, emphasizing shared strategies in coordinating intercellular signaling (Fig. S9D,E). Together, these results position MPOA-projecting, socially activated PIL neurons as key players in orchestrating local signaling during social interactions. Their predicted involvement in multiple communication pathways suggests that they may serve as crucial cellular substrates for mediating behaviorally relevant information flow within the thalamic network.

### Inhibition of PIL function by input from the ventromedial hypothalamic nucleus (VMH)

To identify the upstream factors that allow aggressive behavior to emerge despite the influence of the highly effective aggression suppressing PIL-to-MPOA pathway, we examined the afferent connections of the PIL. Retrograde neuronal tract-tracing using cholera toxin B subunit (CTB) revealed that a large number of VMH neurons project to the PIL, which confirmed previous data (*25*). Aside from cortical areas (medial prefrontal and auditory), the VMH was the only forebrain structure to send significant input to this thalamic region, suggesting a privileged role in modulating PIL activity. We showed that some of the PIL projecting VMH neurons are located in all subdivisions of the VMH. In the region of the ventrolateral subdivision of the VMH where estrogren receptor alpha (ERα) neurons are located, 66.5±6.9% (mean±SD) of CTB neurons were ERα-positive (Fig. 5A-B). Given the well-established role of the VMH and its ERα neurons in promoting aggressive behavior, we hypothesized that this projection could act as a “circuit override,” suppressing the aggression-buffering influence of the PIL-to-MPOA pathway. To test this, we employed a chemogenetic stimulation approach. Adult male rats received AAV injections encoding an excitatory DREADD into the VMH. After allowing time for recovery and viral expression, CNO was microinjected directly into the PIL to selectively activate the PIL terminals of VMH origin (Fig. 5C). Chemogenetic activation of the VMH-to-PIL pathway significantly increased aggressive behavior (Fig. 5D-F). This potentiation of aggression occurred even though the PIL-to-MPOA pathway was intact, indicating that VMH inputs can functionally override its inhibitory influence.

To assess whether this behavioral effect corresponded to suppression of PIL neuronal activity, we measured c-Fos expression following exposure to a male intruder. In control conditions, the conspecific induced robust c-Fos activation in the PIL as expected. Strikingly, chemogenetic stimulation of VMH terminals in the PIL markedly reduced this c-Fos response (Fig. 5G-J), demonstrating that VMH activity can suppress PIL neuronal activation.

These findings suggest a competitive interaction between aggression-promoting VMH inputs and aggression-suppressing tactile pathways converging on the PIL. The VMH-to-PIL projection appears capable of shifting the balance toward aggression by dampening PIL activity, thereby reducing the inhibitory drive transmitted to the MPOA.

## Discussion

The present study identifies the PIL as a previously unrecognized node in the neural circuitry that suppresses aggression. PIL neurons are selectively activated by direct social touch and, through their projections to the MPOA form a thalamic–hypothalamic pathway that functions as an “anti-aggression” circuit (Fig. 6). While the activation of the PIL-to-MPOA pathway suppresses aggression, deprivation of direct physical contact during social isolation, despite preservation of other modalities, leads to heightened aggression comparable to that observed after complete isolation. This indicates that tactile social input is indispensable for maintaining low aggression levels.

**Fig. 6.**
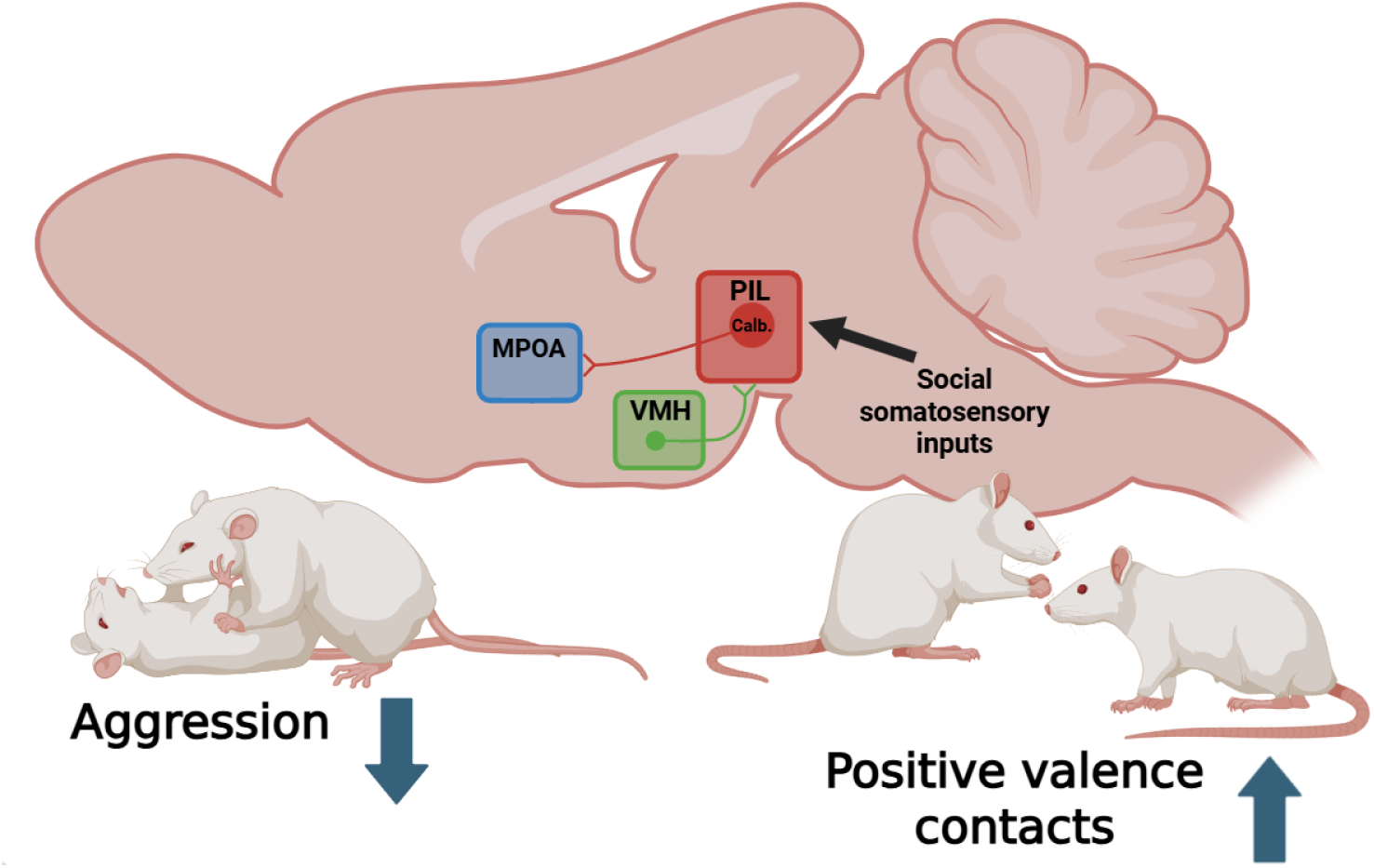
Synopsis of the neuronal pathways reducing aggression in response to social touch. Social touch was demonstrated to activate the PIL neurons, which in turn project to the MPOA, thereby reducing aggression. In contrast, VMH has the capacity to counteract the prosocial effect of the PIL-to-MPOA pathway by inhibiting the PIL. Abbreviations: Calb. – calbindin-positive neuronal cluster, MPOA – medial preoptic area, PIL - posterior intralaminar thalamic nucleus, SN - substantia nigra, VMH – ventromedial hypothalamic nucleus.

### The role of PIL in reducing male aggression

The reduction of aggressive behavior could be of significance in the termination of a fighting bout, as it would prevent injuries that would otherwise be evolutionary disadvantageous. Consequently, the development of neural systems was driven by the need to regulate and control aggressive behavior. While aggression in rodents is primarily driven by the VMH, there are a number of brain regions, such as the medial prefrontal cortex (*26*), the lateral septum (*27*), the bed nucleus of the stria terminalis (*28*), and the medial amygdala (*29*), which are able to reduce aggressive behaviors, primarily by the inhibition of VMH neurons. The present finding that tactile inputs from a conspecific (social touch) are able to reduce aggression is not surprising because fighting itself in the animal kingdom typically involves direct contact between the aggressor and the victim. What could be more efficacious in the prevention of escalation in aggression between two animals than the provision of tactile inputs from the conspecific, thereby engendering a reduction in aggressive motivations? The abrupt termination of attacks in response to optogenetic stimulation of the PIL indicates the involvement of this brain region in the cessation of fighting bouts.

In addition to its role in terminating aggression, social touch may also contribute to the prevention of aggression, which is not warranted during encounters characterized by affiliative social touches. For instance, mating encompasses sexual arousal; however, the degree of concomitant aggressiveness must be meticulously calibrated for successful mating without actually causing harm to the partner.

Besides preventing the escalation of aggression, inputs from conspecifics may reduce aggressive drive, thereby aiding in the evasion of future aggression. These mechanisms are advantageous from an evolutionary perspective as well. In turn, the absence of social touch in animals led to an escalation in aggression, suggesting that social touch plays a critical role in preventing the development of aggression. This suggests that in pair- and group-housed animals, direct social interaction is the modality through which aggression is maintained at low levels within the cage. The observation that optogenetic activation of the PIL not only terminated the ongoing attack but also postponed the onset of the subsequent attack suggests a prolonged effect of PIL neurons. The long-lasting anti-aggressive effect of the PIL neurons is also supported by the reduction of aggressive behavior in the presence of their chemogenetic stimulation. Conversely, the inhibition of PIL neurons augmented the aggressive behavior of rats suggesting that PIL neurons maintain basal activity, the inhibition of which is necessary for the expression of physiological aggression.

From an ecological perspective, it is logical to have mechanisms that allow adaptive aggressions when warranted. The new results provide insights into how the elimination of inhibition by the PIL may take place. The inhibitory influence of PIL neurons on aggression can be overridden by input from the VMH, which partially arrives from ERα neurons of the VMHvl but also from other cell types of the VMH (*30*). The VMH, a well-established area for promoting aggression (*2–4*), projects robustly to the PIL. The chemogenetic stimulation of the VMH-to-PIL pathway not only increased aggression but also suppressed male intruder-induced c-Fos expression in the PIL, indicating a direct inhibitory effect on PIL neuronal activity. This suggests that under conditions where aggression is adaptive, such as territorial defense or competition, the VMH can actively suppress the PIL’s anti-aggression output, allowing the expression of attack behaviors. Since VMH projection neurons are mostly excitatory cells (*31, 32*), they may impose the inhibitory action on PIL neurons via local inhibitory interneurons, which were identified by the present sn-RNAseq study.

The inhibitory action of the PIL on aggressive behavior does not fully explain how it contributes to the long-term reduction of aggression by social contact with conspecifics. The behavioral differences in aggression observed between animals housed in groups and those experiencing social isolation must encompass neuroplastic alterations, given that their behavioral responses to identical stimuli diverge after approximately two weeks. The PIL, in its capacity as a relay center of social touch information for aggression, is poised to contribute to the emergence of neuroplastic changes that have the potential to alter aggressive behaviors. The nature of these neuroplastic changes, however, remains to be determined. Such alterations may encompass gene expression changes or morphological modifications within the implicated neuronal circuits. The candidate brain regions that are potentially altered by neuroplastic changes include the PIL itself, the MPOA, and/or the VMH. Future research may explore this topic.

### The neural systems activating PIL neurons

The fiber photometric results of the present study suggest that social contact activates PIL neurons projecting to the MPOA. This finding extends previous studies on the activation of glutamatergic neurons by social interaction (*16*). The elevated z-score of fiber photometry signal, which commences concurrently with the onset of social touch, indicates that social touch elicits and does not simply follow the activation of PIL neurons. Furthermore, sniffing of the partner animal is not sufficient to activate PIL neurons, thereby corroborating the notion that direct physical contact is requisite for this process. At this juncture, it is worthwhile to engage in speculation regarding which anatomical pathway may be responsible for the activation of PIL neurons. Retrograde tracing was utilized to ascertain the origin of input to the PIL from the cuneate and gracile nuclei, the spinal cord dorsal horn, and the lateral parabrachial nuclei (LPB) (*25*). The somatosensory sensation from a conspecific could arise from the cuneate and gracile nuclei as part of the dorsal column pathway. This pathway has the potential to provide information from fast-conducting, low-threshold mechanoreceptors innervating the skin. However, it may also convey information from postsynaptic dorsal column neurons within the spinal cord dorsal horn. These neurons are capable of transmitting vibrotactile stimuli with high temporal precision, thereby encoding the onset of touch and the intensity of sustained contact (*33*). Alternatively, the anterolateral system may deliver the somatosensory input from the spinal cord to the PIL (*34*). As an alternative option, the social somatosensory input to the PIL might come from the C-tactile peripheral sensory fibers, which are carrying information on gentle stroke (*35*). Within the spinal cord dorsal horn, neurons marked by the G-protein-coupled receptor Gpr83 send axons to the PIL and also to the LPB (*36*), which itself provides input to the PIL (*25*). Gpr83⁺ cells are contacted by mechanosensory afferents expressing Mrgprb4, cells likely corresponding to the C-tactile afferents dorsal root ganglion neurons (*37*). Gpr83+ neurons are also innervated by prokineticin receptor 2-positive spinal interneurons, which engage the brain’s reward circuitry and are linked to affective touch (*38*). Given the reward value of social interactions (*39*), projections of Gpr83+ neurons to the PIL may carry socially relevant tactile signals exchanged between interacting animals.

### Cell types of the PIL involved in the control of aggression

Since local MPOA injection of CNO was as potent in altering aggression as chemogenetic manipulation of PIL, it is likely that PIL neurons projecting to the MPOA mediate this action. A potential target of these projections within the MPOA are ERα-expressing neurons whose projection to the VMH could reduce aggressive behavior (*10*). However, other cell types, e.g. those being activated by social encounter (*8*) cannot be excluded either.

A subsequent investigation of the PIL using sn-RNAseq analysis revealed a population of neurons in the PIL that exhibited significant heterogeneity. A combination of histological and gene expression approaches was used to present evidence that calbindin neurons in the medial portion of the PIL are responsible for reducing aggression by their projections to the MPOA. A substantial augmentation in the quantity of neurons manifesting c-Fos mRNA was identified within these clusters, thereby indicating their participation in the processing of social context. In the event that two groups of socially activated PIL calbindin neurons projecting to the MPOA exhibit distinct gene expression, it is hypothesized that these neurons may also possess different functions, which remain to be determined. The present study identified genes selectively expressed in these clusters, paving the way for future studies to determine their specific functions. The specific functions of the neuropeptides they express, substance P and pituitary adenylate cyclase activating polypeptide (PACAP), respectively, are not yet fully elucidated. While the present study determined the role of the PIL-to-MPOA projections in the reduction of aggression, it also showed that a diverse array of targets of PIL neurons exist raising the possibility that distinct populations of PIL neurons reach other PIL targets, beyond the MPOA. Among the neuronal cell types identified by means of clustering, there could be several projection neurons. We identified two additional clusters of calbindin neurons in the lateral PIL that may not project to the MPOA, but are activated by social encounters. One of these cell groups contains corticotropin-releasing hormone (CRH). To the best of our knowledge, there is an absence of available information regarding their projections and functions. However, given their position overlapping with the medial nucleus of the medial geniculate body, these neurons may be responsive to auditory cues originating from vocalizations. The other cell group containing calcitonin gene-related peptide (CGRP), which has been shown to project to the amygdala (*40*). These neurons may play a role in fear conditioning. Auditory fear conditioning has been demonstrated to occur within PIL neurons (*15*) and their activation by social interaction suggests a potential role in social fear processing.

The presence of several different types of neurons in the PIL suggests that complex information processing occurs in the nucleus rather than simple information relay. It has been proposed that the salience of information is processed here based on multimodal input (*11*). Regarding the interaction between PIL neuronal cell types, cell-cell communication (ccc) analysis revealed that MPOA-projecting, socially activated neurons may receive input from other PIL clusters via semaphorin signaling. Such complex connections between PIL cell types could contribute to neuroplastic changes required for long-term adaptations.

### Translational implications of reducing aggression by social touch

Aggression is a core symptom of many clinical conditions, including dementia, conduct disorder, disruptive/impulse-control disorders, antisocial personality disorder, and autism spectrum disorder. It is also a frequent and dangerous symptom of post-traumatic stress disorder (PTSD), traumatic brain injury, and certain psychotic disorders. While these conditions have different causes, they commonly produce impulsive, reactive, or premeditated aggressive behaviors that cause harm and are difficult to treat. The present data suggest a two-pronged translational approach. Identifying the molecular identity of PIL neuronal clusters could help determine drug targets that would either enhance the PIL-to-MPOA anti-aggression pathway or block VMH-mediated suppression. Alternatively, the results encourage developing structured tactile interventions, such as gentle stroking protocols, caregiver training in affective touch, and dance or movement therapies that include social touch components, as adjuncts to existing treatment programs (*7*). Massage and structured touch can reduce short-term agitation in people with dementia and lower aggressive and disruptive behaviors when administered by caregivers or therapists (*41*). Similarly, dance movement therapy and other embodied interventions can reduce agitation, anxiety, and some behavioral symptoms in people living with dementia and improve affect and social engagement (*42*). These clinical benefits are consistent with experimental data showing that gentle, affective touch has calming, prosocial, and stress-buffering effects in mammals.

In conclusion, the present work establishes the PIL as the first brain region shown to inhibit aggression in response to social touch. The study also defines the PIL’s cellular and projection-specific organization and demonstrates how aggression-promoting hypothalamic inputs can override its function. These findings advance the basic neuroscience of aggression control and lay the groundwork for novel, circuit-based therapeutic strategies aimed at restoring the balance between social affiliation and aggression in clinical populations.

## Materials and methods

### Animals

A total of 126 animals were used, of which 123 were male Wistar rats, and 3 male CD1 mice. The Workplace Animal Welfare Committee of the National Scientific Ethical Committee on Animal Experimentation at Semmelweis and Eötvös Loránd Universities, Budapest, specifically approved this study (PE/EA/117-8/2018, PE/EA/568-7/2021, PE/EA/926-7/2021 and PE/EA/00086/2025, respectively). Thus, the procedures involving rats and mice were carried out according to experimental protocols that meet the guidelines of the Animal Hygiene and Food Control Department, Ministry of Agriculture, Hungary (40/2013), which is in accordance with EU Directive 2010/63/EU for animal experiments. Mice and rats were maintained on a 12h light/dark cycle with lights on at 7am at 22 ± 1°C. Food and water were available *ad libitum.* Paper rolls were provided to the animals as environmental enrichment.

### Stereotaxic surgeries

#### Non-specific viral infection of PIL neurons

For chemogenetic manipulation of PIL neurons, 100 nl of AAV5-hSyn-hM3D(Gq)-mCherry, AAV5-hSyn-hM4D(Gi)-mCherry, or AAV5-hSyn-mCherry was injected into the PIL. The viral vectors were injected bilaterally into the PIL of head-fixed animals. The coordinates were measured from the bregma as follows: anteroposterior (AP) = -5.2 mm, mediolateral (ML) = ± 2.6 mm, and dorsoventral (DV) = -6.8 mm from the dura mater. Subsequent to the surgical procedure, the animals received an injection of enrofloxacin (5 mg/bwkg) to prevent any potential infection and meloxicam (1 mg/bwkg) to reduce pain associated with the surgical procedure. Subsequently, the subjects were separated for a period of three weeks to facilitate their recovery.

#### Viral infection of PIL neurons and optic fiber placement in mice

For optogenetic stimulation of PIL neurons, 75 nl of AAV5-hSyn-hChr2-EYP, was injected into the PIL. The viral vectors were injected unilaterally into the PIL of head-fixed CD1 mice. The coordinates were measured from the bregma as follows: anteroposterior (AP) = -3.05 mm, mediolateral (ML) = +1.5 mm, and dorsoventral (DV) = -3.55 mm from the dura mater. The subsequent step involved the implantation of a fiber optic cannula (RWD fiber optic cannula with a 200 µm core) slightly above the same injection site. After the surgical procedure, the animals received an injection of enrofloxacin (5 mg/bwkg) to prevent any potential infection and meloxicam (1 mg/bwkg) to reduce pain associated with the surgical procedure.

#### vGATE system injection in the PIL and the vGATE protocol

To achieve activity-dependent manipulation of PIL neurons, a novel system was employed, known as vGATE (Virus-delivered Genetic Activity-induced Tagging of Cell Ensembles). This system was employed for the activity-dependent tagging of PIL neurons. A 300-nanoliter (nl) viral cocktail containing rAAV-(tetO)7-Pfos-rtTA; rAAV-Ptetbi-Cre/YC3.60; and rAAV-PhSYN-DIO-hM3D(Gq)-mCherry or rAAV-PhSYN-DIO-hM4D(Gi)-mCherry was injected into the PIL. Following a recovery period of 10 days, during which the animals were single housed, doxycycline (5 mg/kg bw) was injected intraperitoneally. On the subsequent day, the subjects were exposed to a 30-min male intruder test, wherein they were presented with a social stimulus. The resulting social interaction prompted the expression of c-Fos in PIL neurons. The infected neurons concurrently expressed reverse tetracycline transactivator (rtTA). In the presence of doxycycline, rtTA was bound to the promoter region of the second virus (Ptetbi, a bidirectional tetracycline-responsive promoter), thereby enabling CRE-recombinase to be expressed. The third virus was an AAV Cre-dependently expressing inhibitory or stimulatory DREADD. In the course of the study, neurons that demonstrated activity during social interaction were tagged with DREADDs, which were then subjected to subsequent manipulation.

Subsequent to a 10-day period of social isolation, during which the DREADDs were expressed in adequate quantities, behavioral tests were conducted on the animals employing chemogenetic manipulation. The social behavior by the animals that expressed stimulatory and inhibitory DREADDs during the induction of the vGATE system by doxycycline injection was analyzed as described below for chemogenetics.

#### Viral infection of PIL neurons and intracerebral cannula implantation above the MPOA

In order to achieve selective manipulation of the PIL neurons’ axon terminals in the MPOA, 100 nl of stimulatory or inhibitory DREADD-encoding viral vectors (AAV5-hSyn-hM3D(Gq)-mCherry, AAV5-hSyn-hM4D(Gi)-mCherry) were bilaterally injected into the PIL (AP: -5.2 mm, ML: ±2.6 mm, measured from the bregma, and DV: -6.8 mm from the dura mater). Subsequently, intracerebral cannulas were implanted bilaterally above the MPOA (AP: -0.5 mm, ML: ±3 mm, DV: -7.2 mm) at a 16.5-degree angle. The cannulas were affixed to the skull with DURACRYL, a self-curing dental resin.

#### Retrograde viral injection and fiber cannula implantation

Male Wistar rats were anesthetized, and the MPOA (AP: -0.5 mm, ML: -0.6 mm, and DV: -6.7 mm and -7.7 mm) was targeted on one side with 100 nl of a retrogradely spreading virus (AAV-EF1a-jGCaMP8m-WPRE) at both levels. The subsequent step involved the implantation of a fiber optic cannula (RWD fiber optic cannula with a 400 µm core) above the PIL (AP: -5.2 mm, ML: - 2.6 mm, DV: -6.8 mm). This procedure was performed on the same side as the virus injection in the MPOA. The cannulas were affixed to the skull with DURACRYL cement.

#### Retrograde viral injection of MPOA and PIL neurons

The MPOA and the PIL were targeted by the retrogradely transporting cholera toxin B subunit (CTB) in order to examine the input of neuronal cell types in the PIL and VMH, respectively. Nanoinjections were accomplished with glass capillaries (1.14 mm diameter, World Precision Instruments) filled with mineral oil (Sigma-Aldrich), and then 1 µl of CTB was taken up. The glass capillaries were then meticulously positioned on the Nanoinjector. The rats were positioned within a stereotaxic apparatus, with the incisor bar set at a height of −3.3 mm. A 2 mm diameter hole was drilled into the skull, with the drilling being conducted above the target stereotaxic coordinates. The stereotaxic coordinates were determined based on the Paxinos and Watson rat brain atlas (Paxinos and Watson 2006), for the MPOA: AP: − 0.48 mm to the bregma, LM: − 0.6 mm to the midline, DV: − 7.8 mm to the surface of the dura mater; for the PIL: AP = − 5.2 mm to the bregma, ML = + 2.6 mm to the midline, DV = − 6.8 mm to the surface of the dura mater. A total of 100 nanolitres of CTB was injected into the target areas at a rate of 50 nanolitres per minute. The capillary was left in place for five minutes and then withdrawn slowly. Subsequent to the surgical procedures, the animals were permitted a recovery period of two weeks, after which they were euthanised for the purpose of histological analyses.

#### Viral infection of VMH neurons and intracerebral cannula implantation above the PIL

In order to achieve selective manipulation of VMH neurons’ axon terminals in the PIL, 100nl of stimulatory or inhibitory DREADD-encoding viral vectors (AAV5-hSyn-hM3D(Gq)-mCherry) were bilaterally injected into the VMH (AP: -2.86 mm, ML: ±0.8 mm, measured from the bregma, and DV: -9.3 mm from the dura mater). Subsequently, intracerebral cannulas were implanted bilaterally above the PIL (AP: -5.2 mm, ML: ±3.5 mm, DV: -6.94 mm) at a 7.5-degree angle. The cannulas were affixed to the skull using DURACRYL cement.

### Histological approaches

#### Perfusion and tissue collecting

The animals were deeply anesthetized and transcardially perfused with 150 ml of saline, followed by 300 ml of ice-cold 4% paraformaldehyde (PFA) prepared in 0.1 M phosphate buffer at pH=7.4 (PB) for rats and with 15 ml of saline followed by 30 ml of ice-cold 4% PFA prepared in 0.1 M PB for mice. Brains were extracted and postfixed in 4% paraformaldehyde for 24 hours, after which they were transferred to a solution of phosphate-buffered saline (PBS) containing 20% sucrose for a period of two days. Serial coronal sections were obtained by means of a sliding microtome, with each section measuring 40 μm. The sections were then collected in phosphate-buffered saline (PBS) containing 0.05% sodium azide and stored at 4 °C.

#### Immunohistochemistry, antibodies, and cell counting

Subsequent to transcardial perfusion, the brains were sectioned into 40 μm thick coronal slices. A subsample comprising each fifth section was selected for immunohistochemical analysis. In the case of peroxidase reaction-based immunolabelings, free-floating sections were subjected to a treatment comprising 3% hydrogen peroxide for a duration of 15 minutes. Subsequently, the sections were treated with PB containing 0.5% Triton X-100 and 3% bovine serum albumin for a duration of 1 hour. Subsequently, primary antibodies were applied to the sections for a duration of 12 hours. Subsequently, an incubation in phosphate-buffered saline (PBS) containing secondary antibodies was conducted for a period of 60 minutes. The following procedure was carried out: the sample was left for one hour in a solution of avidin-biotin complex (ABC, 1:500). The verification of viral injection sites and mapping projections was achieved through Ni-DAB visualization using chicken anti-mCherry antibodies at a dilution of 1:1,000.

For the purpose of fluorescent immunohistochemistry, free-floating sections were treated with a blocking and permeabilizing solution containing 10% fetal bovine serum, 1% bovine serum albumin, and 0.5% TRITON-X100, dissolved in a 0.1 M PB (pH 7.2), for a period of one hour. The following primary antibodies were then utilized: for calbindin/c-Fos double immunolabeling, a mouse anti-calbindin antibody at a dilution of 1:2000 (Swant, CB300) and a rabbit anti-c-Fos antibody at a dilution of 1:5000 (Abcam, ab190289; for calbindin/CTB double immunolabeling rabbit anti-calbindin antibody at a dilution of 1:3000 (Swant, CB38) and goat anti-CTB antibody at a dilution of 1:10 000 (List Biological Laboratories, #703); for ERα/CTB double immunolabeling anti rabbit ERα antibody at a dilution of 1:900 (Santa Cruz, sc-542) and goat anti-CTB antibody at a dilution of 1:10 000 (List Biological Laboratories, #703). Following the overnight incubation in solutions containing the primary antibodies, the secondary antibodies were applied at a dilution of 1:500 in 0.1 M PB (pH 7.2) for a period of two hours. The secondary antibodies utilized included donkey anti-mouse Alexa Fluor 488 (Jackson Immunoresearch, 715-545-150), donkey anti-rabbit Alexa Fluor 594 (Jackson Immunoresearch, 711-585-152), and donkey anti-goat Alexa Fluor 488 (Jackson Immunoresearch, 705-545-147). DAPI (Merck, D9542) staining was utilized for nucleic acid staining for a period of 10 minutes at a dilution of 1:8000. The sections were mounted using an anti-fade medium (Aqua PolyMount, Polysciences Inc).

The immunolabeling of c-Fos was accomplished through the utilization of rabbit anti-c-Fos antibodies at a dilution of 1:3,000, followed by the visualization of the labeling through an immunoperoxidase reaction. The quantification of c-Fos data was performed by collecting microscopic images (10x magnification) of brain areas on both sides. The number of c-Fos-positive cells was determined by means of cell counting with ImageJ in predefined square boundaries placed within the anatomical region representing the brain areas were utilized to enumerate the number of c-Fos-immunoreactive (c-Fos-ir) cells. In this process, an algorithm based on intensity, size, and circularity thresholding, was employed for all images to select the Fos-ir nuclei. Subsequently, the number of these nuclei was counted automatically by the program. A set of spots was selected on the basis of their brightness intensity value, which was found to be less than 150. The number of c-Fos-containing cells was determined by quantifying the number of selected spots within a size range of 20 to 100 pixels, with a circularity factor ranging from 0.5 to 1.0. The data were meticulously collected from both sides, subsequently averaged, and subjected to rigorous evaluation using a two-way ANOVA.

#### c-Fos studies

The utilization of c-Fos proved to be instrumental in identifying the target areas of PIL projections, which are known to be activated by inputs from the PIL. The animals, which had previously been injected with adeno-associated virus encoding stimulatory DREADDs and expressed with vGATE (n=8), were divided into two groups: half of the animals were administered CNO, while the other half received vehicle. Subsequently, the animals were isolated for a period of two hours, after which they underwent a transcardial perfusion.

The animals that expressed inhibitory DREADDs in their PIL neurons (n=8) were also divided into two groups: half received CNO and the other half received vehicle intraperitoneally. Subsequent to the administration of CNO or vehicle, a male intruder test was performed on the animals for a duration of 30 minutes. Subsequently, the partner animals were removed, and the subjects were left undisturbed for a further 30 minutes prior to the commencement of the perfusion. Animals that expressed stimulatory DREADDs in the VMH were also subjected to a similar experiment: half of the subjects received an intraperitoneal injection of CNO, while the other half received a vehicle injection as a control. Subsequent to the administration of CNO or vehicle, a male intruder test was performed on the animals for a duration of 30 minutes. Subsequently, the partner animals were removed, and the subjects were left undisturbed for a further 30 minutes prior to the commencement of the perfusion.

#### Microscopy and image processing

The sections were examined using an Olympus BX60 light microscope equipped with fluorescent epi-illumination. The images were captured at 2048X2048 pixel resolution using a SPOTXplorer digital CCD camera (Diagnostic Instruments, Sterling Heights, MI) with a 4-40 X objective. Confocal images were acquired with a Zeiss LSM800 confocal microscope system, utilizing a 10x (Plan-Apochromat 10x /0.45) objective and the ‘Tiles’ option of the Zen blue software for lower magnification images. Higher magnification images were made with 20–40X (Plan-Apochromat 20x /0.8 and EC Plan-Neofluar 40x/NA0.75) objectives at an optical thickness of 6 µm for the enumeration of labeled cell bodies. Adjustments were implemented to the contrast and sharpness of the images by employing the “Levels” and “Sharpness” commands within the Adobe Photoshop CS9.0 software. It is noteworthy that throughout the process, the images maintained their full resolution until the final versions were adjusted to a resolution of 300 dots per inch (dpi).

### Behavioral studies

The mice and rats tested for behavior were 12-16 weeks old. Before the behavioral tests, the animals were kept in the procedure room for one week. During the tests, the procedure room was illuminated at 80 lux. The background sound level was measured at 40–50 dB.

#### Isolation of animals from somatosensory social input

The initial phase of the experiment entailed the assessment of the aggression levels exhibited by male Wistar rats, previously housed 3 rats per cage, using a resident intruder test for 30 minutes. Subsequently, the animals were housed in pairs, completely isolated, or separated by a fixed metal grating that permitted visual, auditory, and olfactory interaction between the subjects but prevented somatosensory social contact. Following a two- and four-week period of the experiment, the animals’ levels of aggression were re-evaluated using a 30-minute resident intruder test.

#### Optogenetics

Subsequent to the establishment of a connection between the optic fiber and the subject, CD1 mice (35-40 g) were exposed to a smaller B6 unfamiliar partner (around 30 g) within the confines of their home cage. Prior to the behavioral tests, the subject animals were isolated for a period of three weeks after the surgical procedure, while the partner animals remained in group housing. Subsequently, a 10-minute period of social interaction ensued, during which the animals were subjected to a 1-second-long, continuous photostimulation of the infected PIL neurons during aggressive behavior. Subsequent to the photostimulation phase, the experiment was replicated on the same animals, this time without the photostimulation component.

#### Clozapine-N-oxide administration

Clozapine-N-oxide (CNO) was utilized as a ligand to manipulate neurons expressing DREADDs. CNO was administered intraperitoneally to the animals (0.3 mg/kg bw CNO, dissolved in 5% DMSO in distilled water, 1 ml/kg bw). For the vehicle, 5% dimethyl sulfoxide (DMSO) dissolved in distilled water (1 ml/kg bw) was utilized. A local CNO injection was performed via the intracerebral cannula using a 333x dilution of the previously mentioned solution, with 100 nl of that solution being injected.

#### Behavior experiment for testing aggression

The measurement of aggression was conducted using the resident-intruder test. Subsequent to a two-week or one-month period of social isolation, the subject animals were subjected to resident-intruder tests within the confines of their home cage (length/width/height: 425/285/190 mm). As was the case with the male intruder tests for the chemogenetic studies, the experiment was conducted over a period of three days. Prior to the initiation of the experimental phase, the subjects were isolated for a period of one month at the age of three months. On the first day of the experiment, a vehicle was administered to the animals, and an unfamiliar male rat of similar size was introduced into the subject’s cage, resulting in an aggressive response. On the subsequent day, CNO was injected to manipulate neuronal activity, and the same test was repeated. Subsequent to a 48-hour period of rest, the control injection was repeated. The test was conducted for a duration of 30 minutes, after which the intruder was removed. The intruders were group-housed animals unfamiliar to the experimental animals. They were utilized once per day and reused on the subsequent day. The thirty minutes of video footage were recorded and subsequently analyzed.

The behavioral patterns exhibited by the subjects were meticulously assessed by a single individual who was blinded to the conditions through a manual evaluation process employing the Solomon coder software (*43*). The assessment encompassed various social behavioral elements, including anogenital sniffing, non-anogenital body sniffing, positive-valence direct contacts (e.g., grooming, friendly mounting, and side-to-side contacts), and aggressive behavior. Aggressive behavioral elements were analyzed as dominant mounting (where the subject pins down the partner), fighting (where the animals tumbled over, kicked, punched, and boxed each other), and biting.

#### Elevated Plus Maze test

A protocol was devised to assess anxiety-like behavior and locomotion. This protocol included the elevated plus maze test (EPM) performed on a day separate from those devoted to testing aggression. Half of the animals in each group (stimulatory, inhibitory, or control virus injected) received CNO for chemogenetic manipulation and were compared with animals that received vehicle. Prior to the experimental procedure, the animals had not been habituated to the apparatus. Ten minutes of footage was recorded, documenting their behavior. Subsequently, the entries were recorded and the time spent in the open or closed arms was also evaluated.

### Fiber photometry experiments

Three weeks after the surgical procedure, the animals demonstrated signs of recovery and were subsequently subjected to behavioral testing. The activity of PIL neurons was measured using an R811 Dual Colour Multichannel Fiber Photometry System (RWD, Shenzen, China). Subsequent to the establishment of a connection between the fiber cannula (with ceramic ferrule, 2.5 mm diameter, 300 µm core with NA) and the photometry system via an optical fiber, the subject animal was positioned within an arena for a duration of one minute. Subsequently, an unfamiliar male Wistar rat was introduced into the arena. The animals were permitted to interact with each other for a period of five minutes, during which the activity of the PIL neurons was recorded. Video documentation of the animals’ behavior was also recorded. The 410 nm light was utilized as an isosbestic light, with the objective of recording noise and artifacts caused by movement. The excitation of GCaMP was achieved through the utilization of 470 nm light at 50 Hz, and the subsequent collection of data pertaining to the activation of PIL neurons was facilitated.

The analysis was conducted using RWD software. Initially, the data underwent a process of smoothing and motion correction, with the isosbestic data being extracted from the recording to ensure the removal of motion artifacts. The activity measured in the PIL was then correlated with the behavioral elements displayed by the animals. The peri-event analyses were initiated two seconds prior to the presentation of the behavioral element, subsequently followed by a six-second recording of the activity of PIL neurons. The physical contact between the animals and the subsequent separation of the subjects (i.e., when the subject was moving away from their partner animal) were analyzed. A total of 47 events involving physical contact and 28 events involving moving away were recorded. Z-scores of deltaF/F were calculated by the software for each event. The area under the curve (AUC) was calculated for the Z-scores during the 2-second period following the presentation of the behavioral elements and the 2 seconds prior to this.

### Single-nucleus sequencing of PIL neurons

#### Experimental design

Four of the animals were isolated, while the remaining eight were paired and housed together for a period of two weeks. On the day prior to the dissection, the animals were separated for a 24-hour period to establish a baseline for c-Fos expression. On the day of the dissection, the 4 aggressive animals and 4 non-aggressive animals underwent a 20-minute resident-intruder test. One hour after the initiation of the experiment, the animals were terminated and the PIL was collected with the micropunch technique. It was then immediately frozen in 2-methyl butane solution on dry ice and then stored at -80°C. The remaining four non-aggressive animals were utilized as controls and did not receive any partner animals prior to the dissection. Their PIL was similarly processed as for the other two groups.

#### Tissue collection and nucleus isolation for RNA sequencing

Frozen brain tissues containing the posterior intralaminar thalamic complex (PIL) were dissected from adult male Wistar rats (n=12), representing three experimental conditions: control (Cont, n=4), affiliative (Affi, n=4), and aggressive (Aggr, n=4) phenotypes. Tissue from the 4 animals of each groups was pooled to result in 50-100 mg of fresh-frozen tissue, which was used for single-nucleus RNA sequencing. Nuclei isolation was performed using a proprietary detergent- and enzyme-based protocol optimized for brain tissue, involving mechanical dissociation, nuclear lysis, and density-based purification steps. All steps were processed by Novogene Europe (Cambridge, UK). Single-nucleus suspensions were generated using a dounce homogenization step, followed by density gradient centrifugation.

#### Library preparation and sequencing

Library preparation was performed by Novogene using the Chromium Next GEM Single Cell 3′ Reagent Kits v3.1 (10x Genomics). Libraries were sequenced on an Illumina NovaSeq 6000 platform to obtain paired-end reads (2 × 150 bp).

#### Preprocessing and quality control

Raw sequencing data were processed with CellRanger v9.0.1 (*44*), using a custom reference genome (mRatBN7.2). The resulting count matrices were imported into Seurat v5.3.1 (*45*) in R (v4.4.3). Raw and filtered UMI count matrices were processed with SoupX v1.6.2 to correct for ambient RNA contamination (*46*). Briefly, for each sample, raw (raw_feature_bc_matrix) and filtered (filtered_feature_bc_matrix) matrices were loaded into SoupChannel objects. Cluster assignments were imported from an initial Seurat object, and contamination fractions were estimated using autoEstCont. The corrected UMI counts were obtained via adjustCounts, and new Seurat objects were created for each sample based on the corrected matrices. Quality control was performed on each corrected object by computing per-cell quality metrics including the number of detected genes (nFeature_RNA), total UMI counts (nCount_RNA), and the percentage of mitochondrial transcripts (percent.mt) (Fig. S5A, Table S1). Cells with fewer than 200 or more than 5,000 detected genes, or with >5% mitochondrial content, were removed from the analysis.

#### Normalization and detection of doublets

Following quality control and ambient RNA correction, SCTransform normalization was performed using Seurat’s SCTransform() function (vst.flavor=“v2”), regressing out technical covariates such as nCount_RNA. Doublets were identified using scDblFinder v 1.18.0 (*47*) on the SCTransform-normalized SingleCellExperiment object, specifying sample identity and including 20 principal components. The predicted doublet class was assigned back to the integrated Seurat object metadata. Clusters with a high proportion of predicted doublets (>70%) were excluded from further analysis. In addition, one cluster showing abnormally high expression of mitochondrial genes and heat-shock proteins (e.g. mt-Rnr1, mt-Rnr2, Hsp90ab1) was removed due to potential cell stress or degradation.

#### Integration and clustering

Samples were split by original identity and normalized independently using SCTransform as described above. The top 2,000 highly variable genes were identified using SelectIntegrationFeatures(). Datasets were prepared for integration using PrepSCTIntegration() and integrated with SCT-based normalization using FindIntegrationAnchors() and IntegrateData(), specifying reciprocal PCA across samples. The resulting integrated dataset was used for all downstream analyses. Principal component analysis (PCA) was performed on the integrated object using 50 dimensions. Uniform Manifold Approximation and Projection (UMAP) was computed using the top 15 principal components with a minimum distance of 0.5. Clustering was carried out using a shared nearest neighbor graph followed by Louvain community detection (FindNeighbors() and FindClusters(), resolution=0.5). The number of informative PCs was determined using elbow plot and cumulative explained variance analysis.

#### Cluster annotation and marker gene analysis

Cluster-specific marker genes were identified using the FindAllMarkers function with the MAST method (*48*). Only genes expressed in at least 10% of cells in the cluster showing a log2 fold change ≥ 0.5 with adjusted p-value <0.05 (Bonferroni correction) were retained. Canonical marker genes for major brain cell types (e.g., Snap25, Rbfox3 for neurons; Slc1a2, Aqp4 for astrocytes; Plp1, Opalin for oligodendrocytes; Pdgfra, Cspg4 for OPCs; Pecam1, Flt1 for endothelial cells; Abcc9, Rgs5 for pericytes; Ptprc, Apbb1ip for microglia; Slc17a6, Slc17a7 for glutamatergic neurons; Gad1, Gad2, Slc32a1 for GABAergic neurons; and Th for dopaminergic neurons) were used to assign cell type identities (Fig. S5C). Annotation was supported by hierarchical clustering and average expression patterns of canonical markers across clusters. Neuropeptide gene expression was evaluated using manually curated gene lists, and cluster-specific expression levels were quantified (mean log2FC, adjusted p-value, and percent of expressing cells) using the Wilcoxon rank-sum test implemented in the FindAllMarkers function.

#### Subclustering of neurons

After subsetting the Seurat object to include only neuronal cells, unsupervised clustering was performed. In the resulting clusters, some showed highly imbalanced sample representation, where at least one of the three biological samples contributed less than 10% of the total cells, while another sample contributed over 80%. This likely resulted from slight anatomical variation in tissue dissection between animals, possibly including adjacent regions in some samples but not others. To ensure robustness, these underrepresented clusters were excluded from further analysis. The remaining cells (clusters with>10% contribution from all samples) were re-clustered, resulting in 18 neuronal subclusters. All downstream analyses including neuropeptide expression, cell-cell communication inference, and quantification of c-Fos expression were performed on this filtered and re-clustered neuron population.

#### Cell-cell communication analysis

Intercellular communication between neuronal subclusters was analyzed using CellChat v1.6.1 (*49*). As no rat-specific ligand–receptor database is available, we adapted CellChatDB.mouse by retaining only interactions involving genes with confirmed rat orthologues, based on the MGI Homology Report (HOM_AllOrganism.rpt.txt) downloaded from the Mouse Genome Informatics (MGI) database (https://www.informatics.jax.org/downloads/reports/). Secreted signaling pathways were analyzed, considering only genes expressed in ≥10% of cells (thresh.pc=0.1) and interactions involving at least 10 cells. Communication probabilities, pathway-level aggregation, and network centrality scores were computed using standard CellChat functions. Signaling roles and global communication patterns were visualized using scatter plots, river plots, and dot plots.

To extract higher-order organizational patterns, we applied non-negative matrix factorization (NMF) to the global communication matrices. This approach grouped clusters into shared incoming and outgoing signaling programs. Distinct incoming and outgoing communication patterns were visualized using both clustered heatmaps and Sankey (alluvial) diagrams. While both reflect the same underlying data, the heatmaps highlight the relative contribution of each cluster to a given pattern, whereas the Sankey plots emphasize flow dynamics between clusters, patterns, and pathways.

### Statistical analysis

The statistical calculations were performed using GraphPad Prism. Initially, the Shapiro-Wilk test was conducted on the entire dataset to ascertain whether the data followed a Gaussian distribution. For normally distributed data, two-tailed paired t-tests or unpaired t-tests were used. In instances where the data did not conform to a normal distribution, the Wilcoxon test (AUC of physical touch) or the Friedman test followed by the Dunn’s post hoc test were employed. In instances where a greater number of variables were compared, a 2-way ANOVA (quantitative analysis of c-Fos positive cells) or a repeated measure 2-way ANOVA (testing aggressive behaviors) was employed. Significant changes are indicated as 0.010 <*p <0.050, 0.001 <**p <0.010, and ***p <0.001.

## Acknowledgments

The technical assistance of Nikolett Hanák and Ágnes Konrád is appreciated. The authors would also like to thank Dr. Ted. B. Usdin and Dr. Sarah K. Williams Avram of the National Institute of Mental Health, Bethesda, for critically reading tha manuscript. During the preparation of this work, the authors used publicly available DeepL Write tool (https://www.deepl.com/write) in order to improve the wording and grammar of the text. After using this tool/service, the authors reviewed and edited the manuscript and take full responsibility for the content of the publication.

## Funding

Hungarian Academy of Sciences grant NAP2022-I-3/2022 (AD)

Hungarian Academy of Sciences grant NAP-KOLL-2023-2/2023 (AD)

Hungarian Academy of Sciences grant NAP-KOLL-2023-6/2023 (AD)

National Research, Developmental and Innovation Office grant NKFIH OTKA K146077 (AD)

National Research, Developmental and Innovation Office grant NKKP Excellence 151425 (AD)

Ministry of Culture and Innovation, in collaboration with the National Research, Development and Innovation Office, grant 2024-1.2.2-ERA_NET-2025-00021 (AD)

Higher Education Institutional Excellence Programme of the Ministry of Human Capacities in Hungary, within the framework of the Neurology Thematic Programme of Semmelweis University, TKP 2021 EGA-25 (AD)

Eotvos Lorand University Central Europe Leuven Strategic Alliance Research Fund CELSA/24/020 (AD)

Strategic Research Fund of the University of Veterinary Medicine Budapest, Grant No. SRF-002 (GP)

New National Excellence Program of the Ministry for Culture and Innovation from the source of the National Research, Development and Innovation Fund, EKÖP-2024-15 (FD) New National Excellence Program of the Ministry for Culture and Innovation from the source of the National Research, Development and Innovation Fund, EKÖP-2024-112 (TL)

New National Excellence Program of the Ministry for Culture and Innovation from the source of the National Research, Development and Innovation Fund, EKÖP-2025-591 (TL)

Synergy European Research Council (ERC) grant “OxytocINspace” 101071777 (VG)

SFB Consortium 1158-3 (VG)

German-Israeli Program (DIP) GR3619-1 (VG)

ERANET-Neuron grant GR 3619/25-1 (VG)

## Author contributions

Conceptualization: DK, AD

Methodology: TL, BBD, DK, FD, IC, VS, GP, VG, AD

Investigation: TL, BBD, DK, FD, IC, GP, AD

Visualization: TL, FD, GP, AD

Funding acquisition: TL, BBD, FD, GP, VG, AD

Project administration: GP, AD

Supervision: DK, AD

Writing – original draft: TL, FD, AD

Writing – review & editing: TL, FD, GP, VG, AD

## Supplementary Materials

### Supplementary figures

**Fig. S1.**
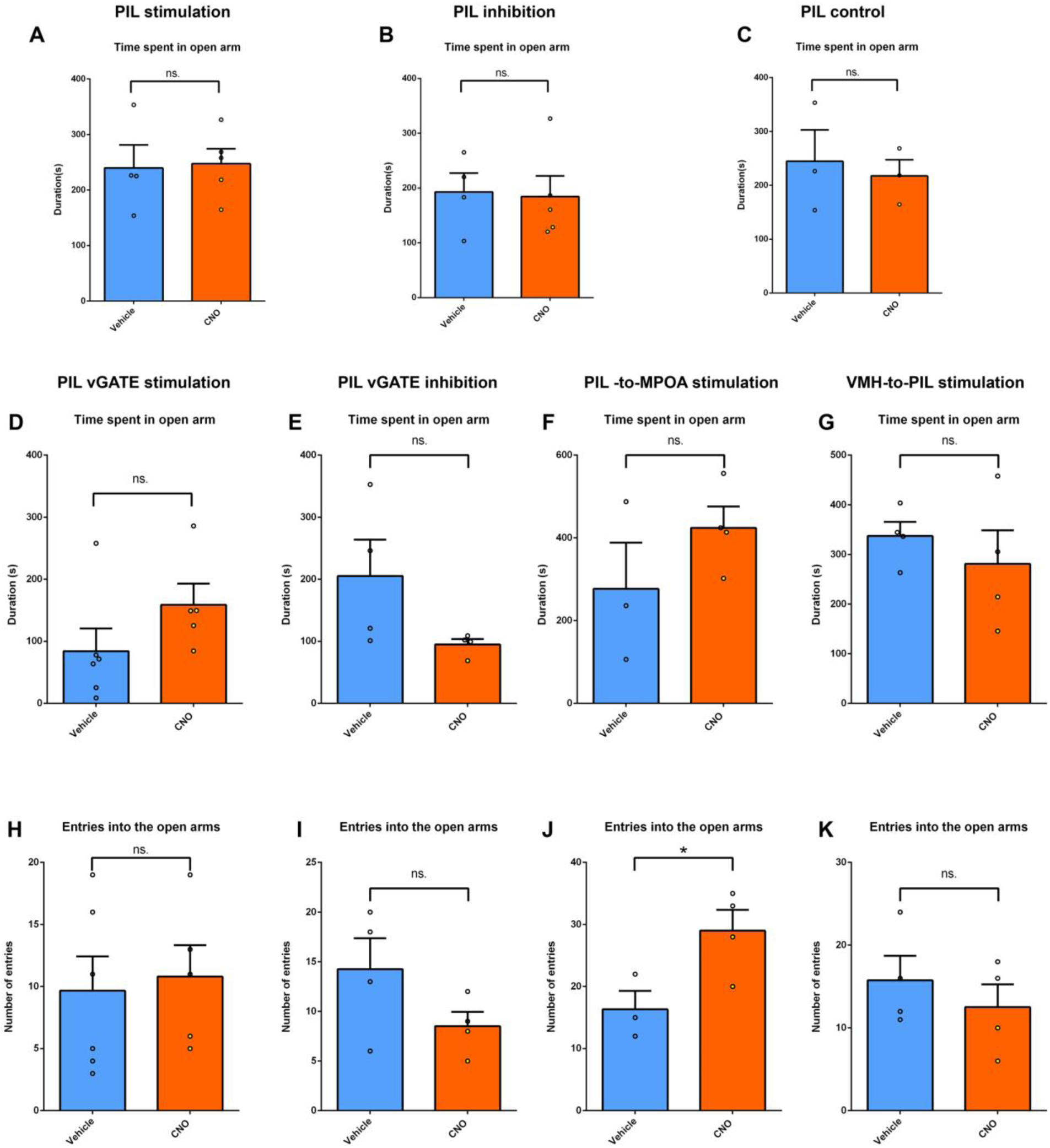
The lack of effect of the PIL neurons on locomotion and anxiety. (**A-C**) The duration that subjects spent in the open arm of the elevated plus maze apparatus was measured during chemogenetic manipulation of PIL neurons. Stimulation, inhibition, and control injections did not influence anxiety-like behavior. (**D-E**) The duration of the subject’s presence in the open arm of the elevated plus maze apparatus was measured during chemogenetic stimulation and inhibition of socially-tagged PIL neurons. Neither stimulation (D) nor the inhibition (E) of socially tagged PIL neurons affected anxiety-like behavior. (H–I) shows the number of entries into the open arm during stimulation (H) and inhibition (I) of socially-tagged PIL neurons. Manipulation of socially tagged PIL neurons had no effect on locomotor activity. (**J-K**) The number of entries into the open arm during stimulation of the PIL-to-MPOA pathway (J) and the VMH-to-PIL pathway (K), which did not affect locomotor activity.

**Fig. S2.**
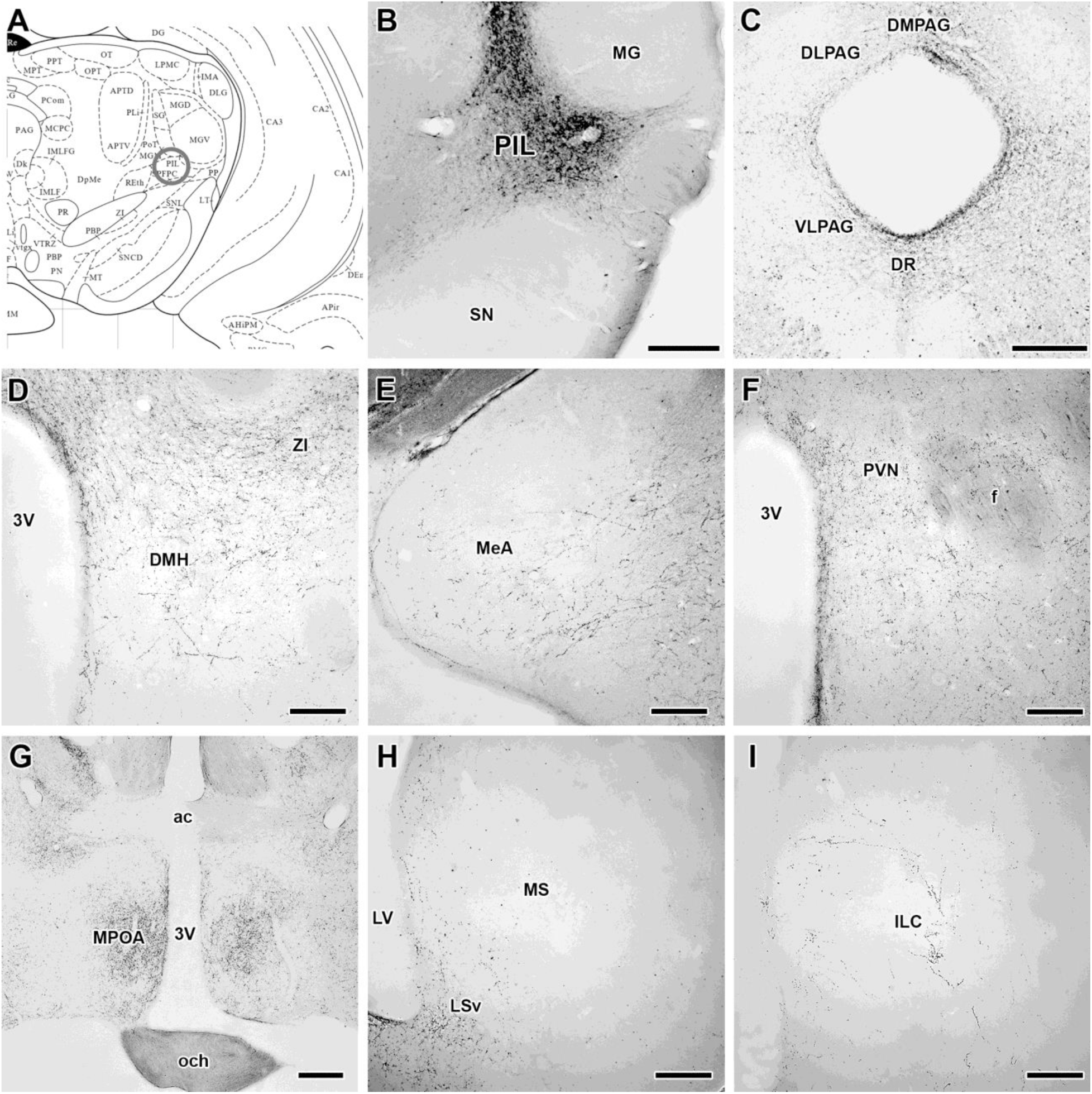
Projections of the socially activated PIL neurons. (**A**) The location of the PIL as indicated in the Paxinos rat brain atlas. (**B**) The injection site of the vGATE viral cocktail into the PIL and tagged socially activated neurons. (**C-I**) The fiber terminals of socially-tagged PIL neurons in the periaqueductal gray (PAG) (C), the dorsomedial hypothalamic nucleus (DMH) (D), the medial amygdaloid nucleus (MeA) (E), the paraventricular hypothalamic nucleus (PVN) (F), the medial preoptic area (MPOA) (G), the ventral subdivision of the lateral septal nucleus (LSv) (H), and the infralimbic cortex (ILC). Scale bar = 500 µm for the PIL, PAG and MPOA; 250 µm for the DMH, MeA, PVN, LS and ILC.

**Fig. S3.**
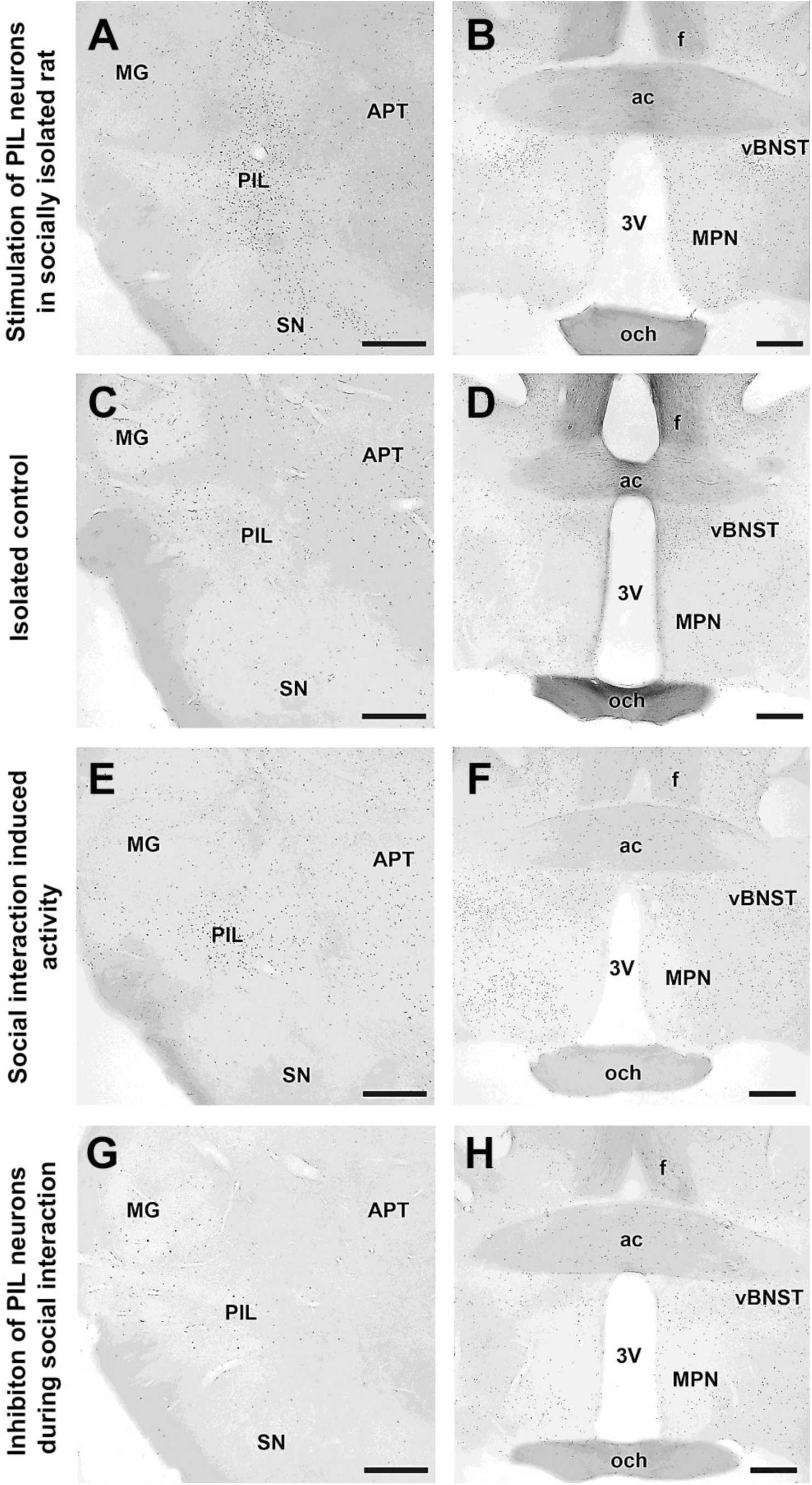
The influence of chemogenetic stimulation and inhibition on c-Fos activation. (**A**) A substantial number of neurons that express the protein c-Fos were identified in the PIL following chemogenetic stimulation of PIL neurons in the absence of social interaction. (**B**) Elevated levels of c-Fos expression in the MPOA were observed in response to chemogenetic stimulation of PIL neurons in the absence of social interaction. (**C**) Control c-Fos expression in PIL neurons following vehicle administration in the absence of social interaction. (**D**) Control c-Fos expression in MPOA neurons following vehicle administration in the absence of social interaction. (**E**) The absence of c-Fos expression in PIL neurons following aggressive interaction and chemogenetic inhibition of the PIL via CNO administration. (**F**) A subsequent reduction in c-Fos expression was observed in MPOA neurons following aggressive interaction and chemogenetic inhibition of the PIL via CNO administration. (**G**) The social interaction-induced c-Fos expression in the PIL was utilized as control. (**H**) The social interaction-induced c-Fos expression in the MPOA was utilized as a control. Scale bar=500 μm in every panel. Abbreviations: ac – anterior commissure, APT – anterior pretectal nucleus, vBNST – bed nucleus of stria terminalis, ventral part, f – fornix, MG – medial geniculate body, och – optic chiasm, PIL - posterior intralaminar thalamic nucleus, SN - substantia nigra, 3V – 3^rd^ ventricle.

**Fig. S4.**
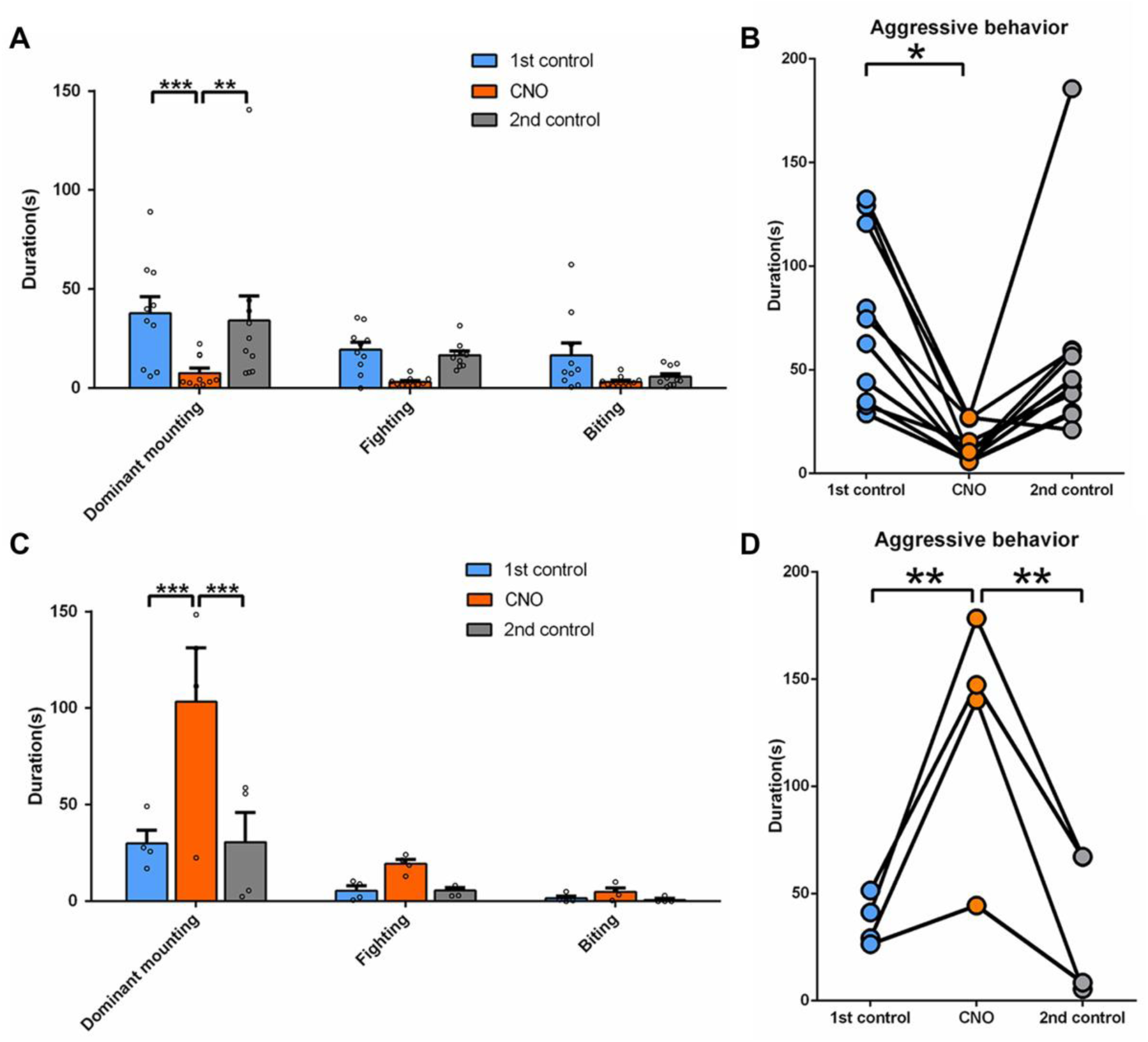
The effect of socially-tagged PIL neurons on aggression. (**A**) Chemogenetic stimulation of socially tagged neurons inhibited dominant mounting but also tended to reduce fighting and biting. (**B**) The sum of these behavioral elements, called aggression, was reduced on the CNO day. (**C**) Chemogenetic inhibition of socially-tagged PIL neurons increased dominant mounting and tended to elevate fighting and siting. (**D**) Aggression was increased in response to CNO treatment as compared to both the previous and the subsequent control days.

**Fig. S5.**
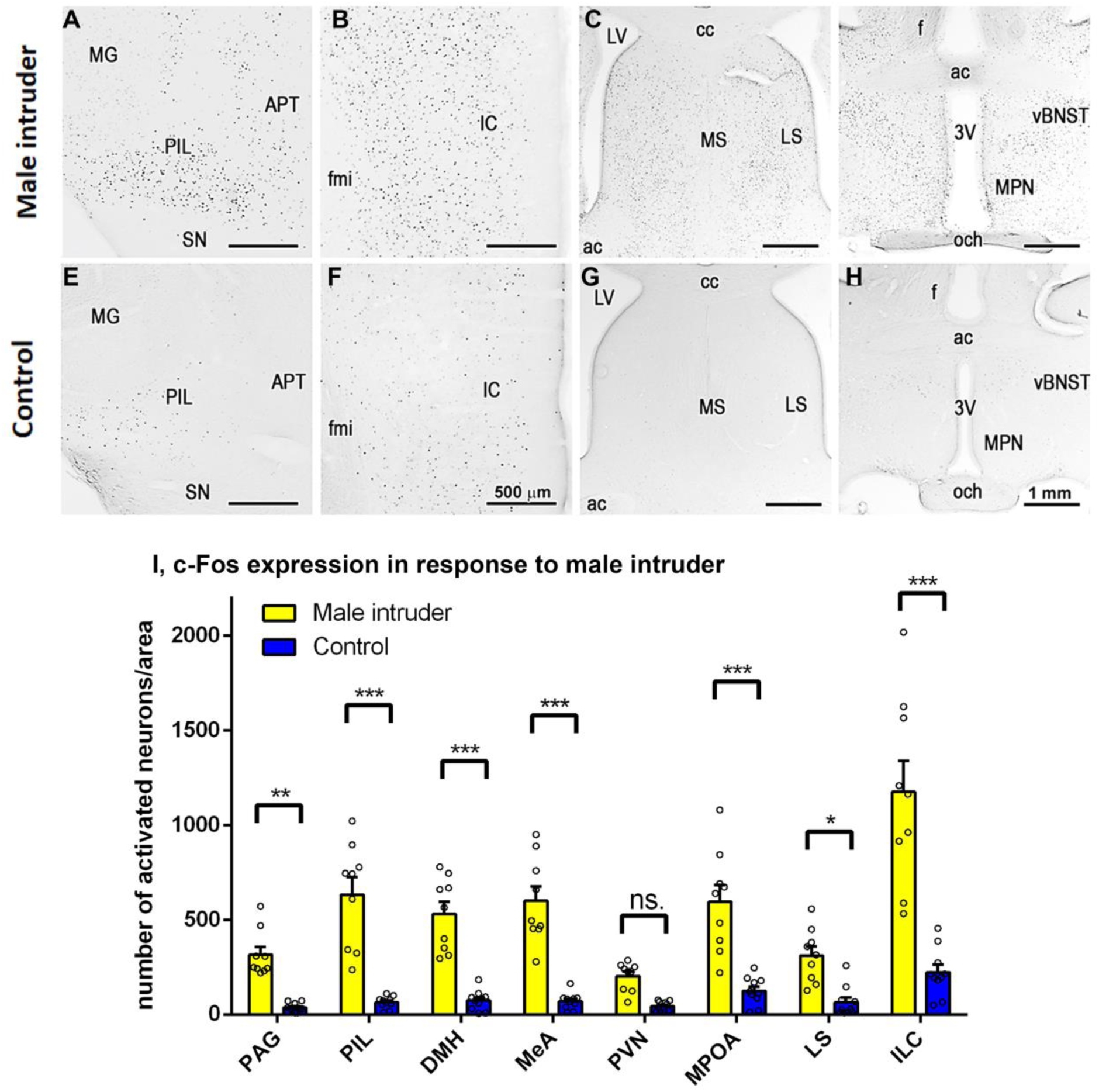
Male intruder-induced activation in the PIL and in forebrain regions projected to by PIL neurons. c-Fos expression induced by aggressive social interaction in a male rat receiving an unfamiliar male into the home cage. (**A**) Posterior intralaminar thalamic nucelus (PIL), (**B**) infralimbic cortex (IL), (**C**) lateral septal nucleus (LS), (**D**) medial preoptic area (MPOA). (**E-H**) Significantly less c-Fos expression is present in isolated control male rats in the same brain regions. (**I**) Quantification of c-Fos expression in response to a male intruder in the PIL and in forebrain regions projected by PIL neurons.

**Fig. S6.**
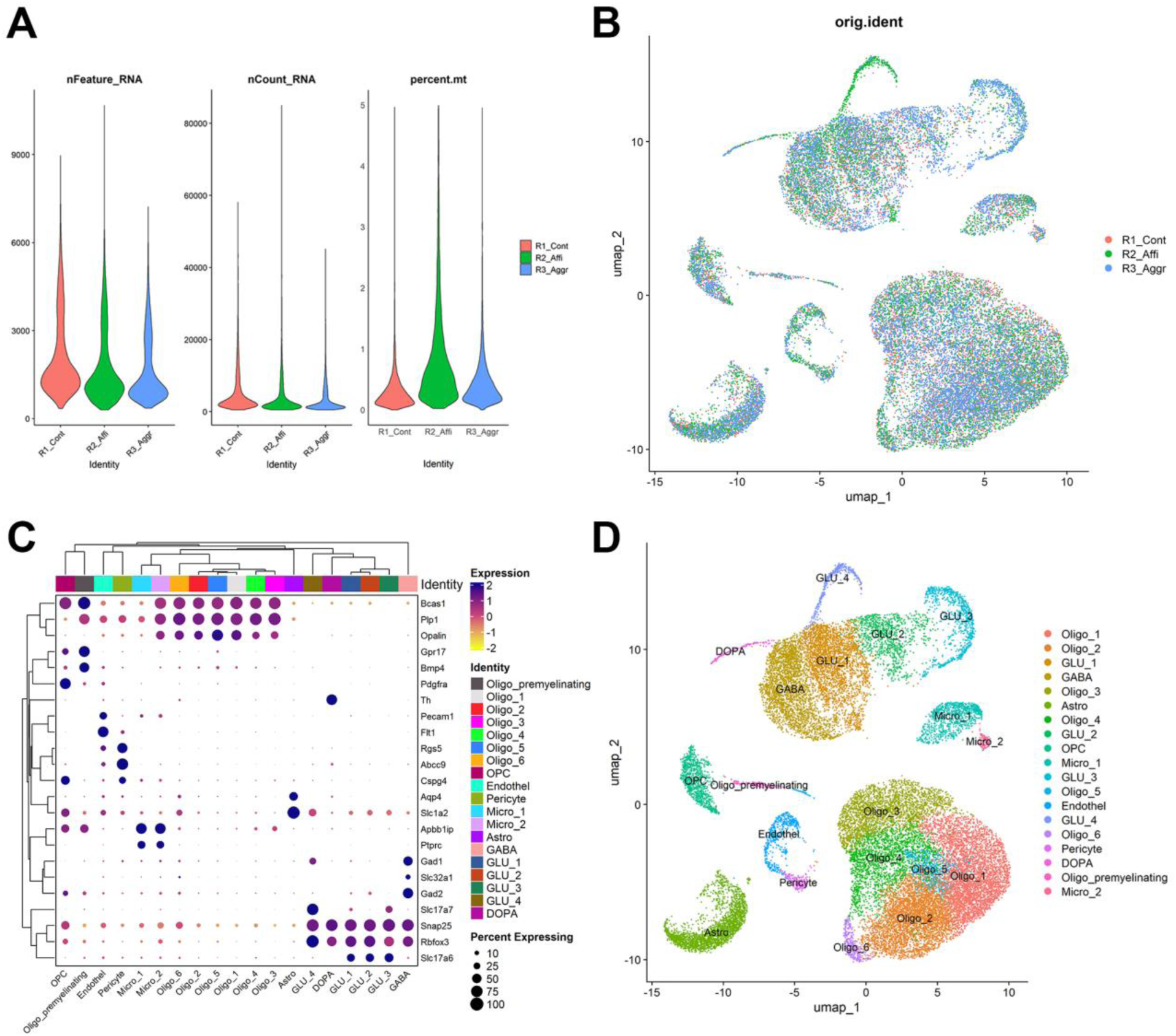
Cellular composition and quality control metrics across pooled PIL samples. (A) Violin plots of gene number (nFeature_RNA), count number (nCount_RNA)), and mitochondrial percentage (percent.mt) in PIL samples. The dataset was filtered out based on these violin plots. (B) Uniform Manifold Approximation and Projection (UMAP) embedding colored by sample origin showing three pooled rat PIL samples: R1_Cont (control), R2_Affi (affiliative), and R3_Aggr (aggressive). (C) Dot plot of canonical marker genes used for manual cell type annotation. Gene expression patterns supported the identification of major brain cell types including glutamatergic neurons, GABAergic neurons, oligodendrocytes, astrocytes, microglia, endothelial cells, pericytes, and ependymal cells. (D) UMAP embedding colored by final cell type annotation, highlighting diverse cell populations derived from pooled PIL samples.

**Fig. S7.**
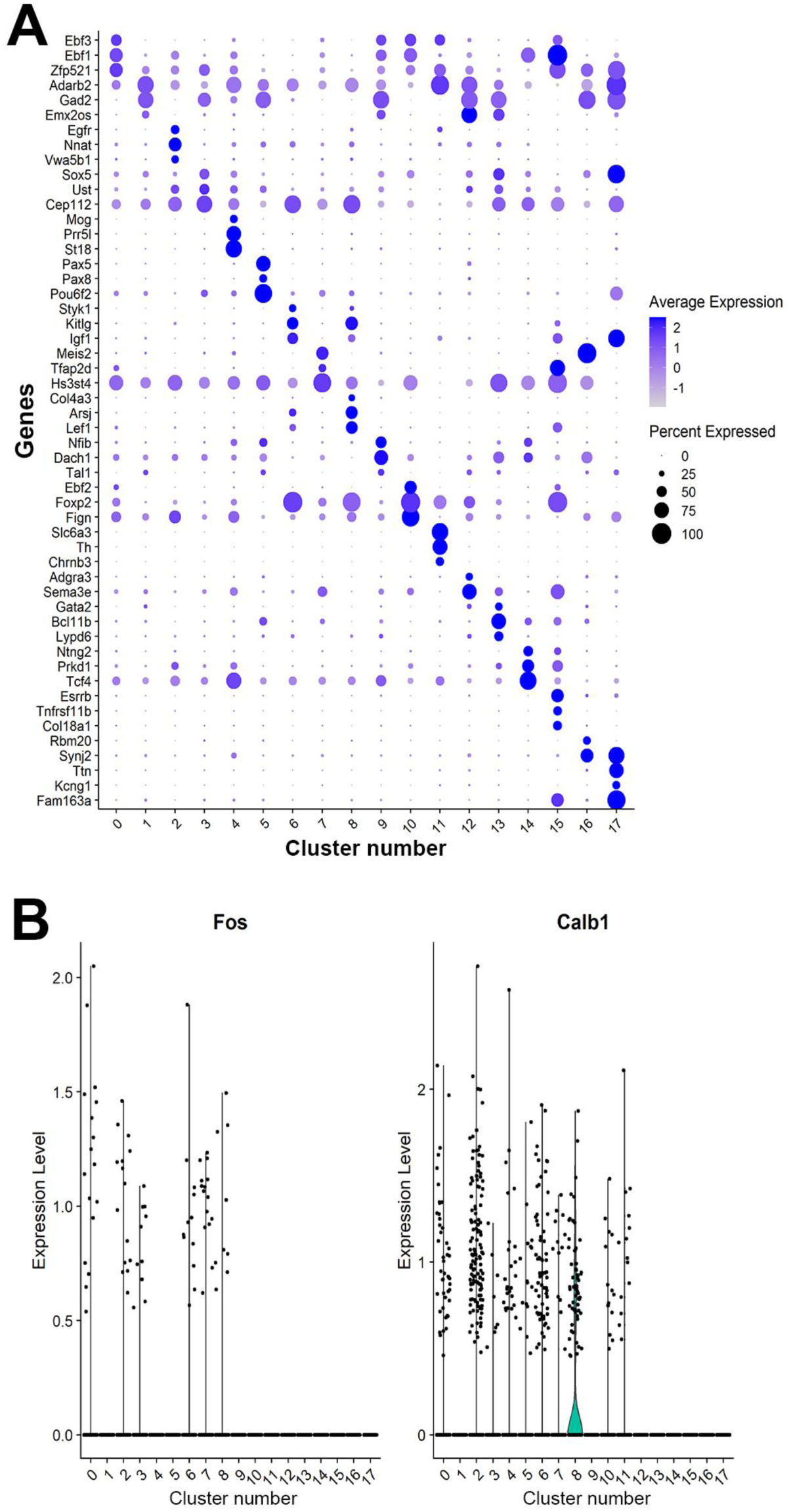
Expression patterns of neuronal subclusters. (A) A dot plot showing the expression features of three signature genes selected from each cell cluster among the neuronal subclusters. Gene expression levels, ranging from low to high, are indicated by a color gradient from gray to blue. The percentages of cells expressing specific genes are proportional to the size of the dots.(B) Violin plots showing the expression levels of c-Fos (left) and Calb1 (right) across all 18 neuronal subclusters.

**Fig. S8.**
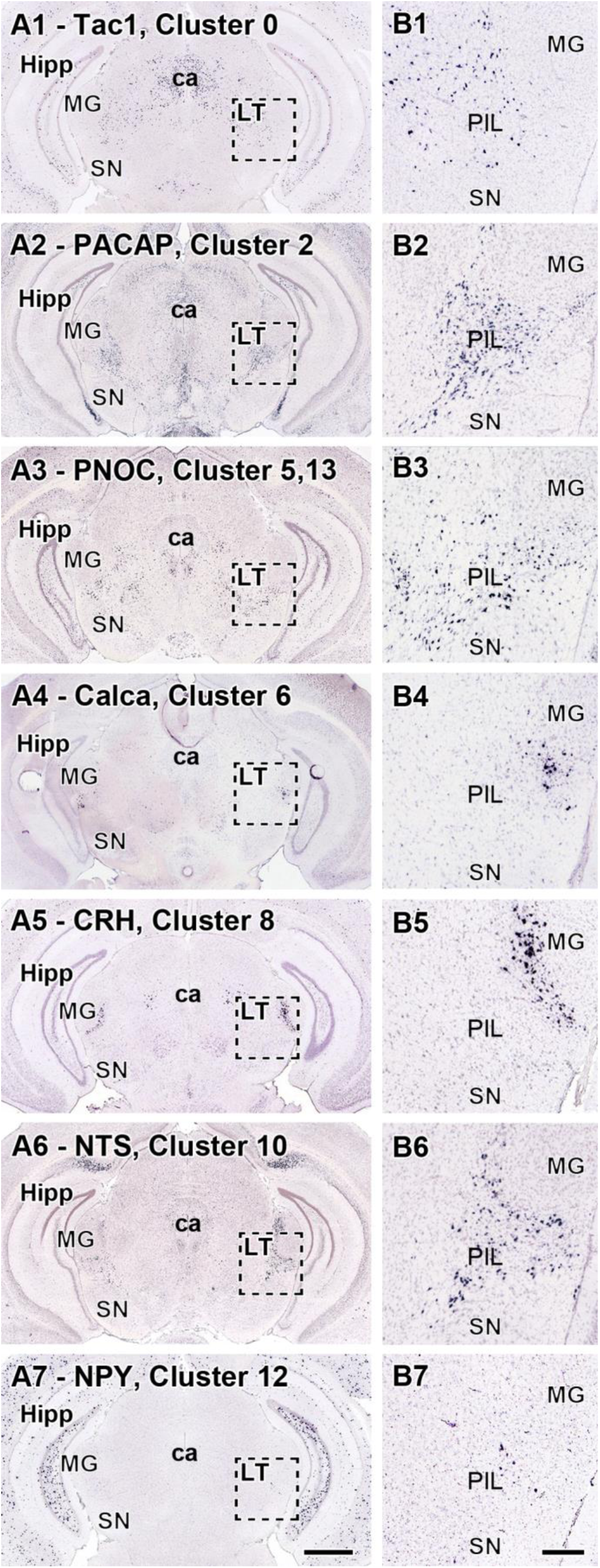
The distribution of PIL neurons expressing neuropeptides characteristic of specific clusters. The panels are from the Allen Brain Atlas. In situ hybridization histochemistry shows the location of neuropeptide mRNA expression in coronal sections at the level of the PIL. (**A1**) Tachykinin 1 (Tac1) encoding substance P, (**A2**) Adcyap1 encoding Pituitary adenylate-activating polypeptide (PACAP), (**A3**) Pronociceptin (PNOC), (**A4**) Calcitonin A (Calca) encoding Calcitonin gene-related peptide (CGRP), (**A5**) Corticotropin-releasing hormone (CRH), (**A6**) Neurotensin (NTS), (**A7**) Neuropeptide Y (NPY). (**B**) High magnification images of the framed areas in the corresponding A panel to show the location and distribution of neuropeptide-expressing neurons in the PIL. Scale bar = 1 mm for A and 200 µm for B. Additional abbreviations: ca – cerebral aqueduct, Hipp – hippocampus, LT – lateral thalamus, MG – medial geniculate body, SN – substantia nigra.

**Fig. S9.**
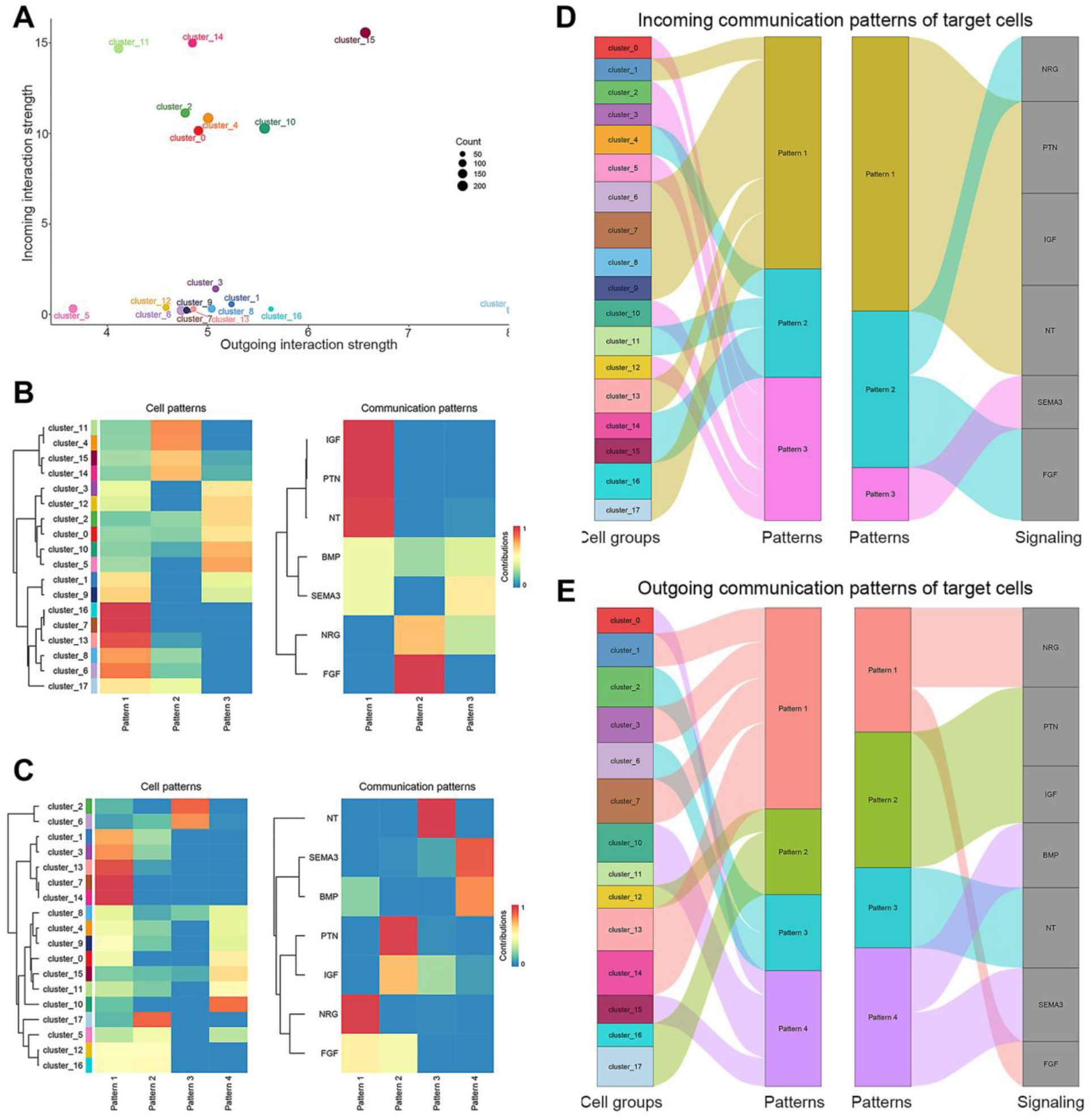
Analysis of cell-cell communication. (A) The strengths of cell-cell communication inferred by CellChat are displayed for each neuronal cluster in PIL samples. The x-axis of the graph corresponds to the outgoing interaction strengths, while the y-axis corresponds to the incoming interaction strengths. The diameter of the circle is indicative of the number of predicted ligand-receptor interactions, thereby providing a quantitative measure of the communication capacity of each cluster. (B) Heatmaps showing the hierarchical clustering of neuronal clusters (left) and signaling pathways (right) based on incoming communication patterns. (C) Heatmaps illustrating clustering based on outgoing communication patterns. (D) Sankey diagram showing incoming cell–cell communication patterns among the neuronal subclusters. Incoming patterns show how target clusters coordinate with each other and with specific signaling pathways to respond to incoming signals. (E) Outgoing signaling patterns of secreting cells, illustrating the correspondence between inferred latent patterns and cell groups, as well as signaling pathways. The thickness of the flows indicates the contribution of each cell group or signaling pathway to the respective latent pattern, while the height of each pattern reflects the number of associated cell groups or pathways. The analysis of outgoing patterns reveals the coordination mechanisms between sender cells and the specific pathways involved in driving intercellular communication.

### Supplementary Table Interpretations

**Table S1. Quality control metrics for single-nucleus RNA-seq libraries.** This table summarizes the following key quality metrics for each sample, including the estimated number of nuclei, mean reads per cell, median genes and UMIs per cell, total reads, and barcode/UMI validity rates.

**Table S2. Cluster-specific marker genes for major cell types in the PIL.** This table lists the marker genes identified for each major cell type cluster. The genes are ranked by adjusted p-value, log₂ fold change, and percent expression within the cluster. These marker genes were determined using differential expression analysis, which enables robust annotation of cell type identity and delineation of major non-neuronal and neuronal populations within the PIL.

**Table S3. Differential expression analysis of neuronal subclusters.** This table presents the results of differential gene expression analysis for each of the 18 neuronal subclusters. This information provides the molecular basis for subcluster annotation and the identification of distinct neuronal phenotypes within the PIL.

**Table S4. Neuropeptide expression across neuronal subclusters.** This table summarizes the expression of key neuropeptide genes across neuronal subclusters. For each neuropeptide, the table lists the average log2 fold change, adjusted p-value, and the proportion of cells expressing the gene within each cluster. These data highlight the diversity and specificity of neuropeptide signaling within the PIL neuronal landscape.

**Table S5 c-Fos and Calb1 expression across neuronal subclusters and samples.** This table quantifies the number of c-Fos- and Calb1-positive cells in each subcluster, broken down by sample. For each subcluster and sample, it shows the number and proportion of c-Fos⁺ and Calb1⁺ cells, as well as the total number of cells per subcluster. These data provide insight into the activation status and molecular heterogeneity of specific neuronal subpopulations under different experimental conditions.

**Table S6.** Statistical details of experiments.

| Related Figure | Sample size | Mean + SEM | Statistical test and p-values |
| --- | --- | --- | --- |
| Fig. 1 A | Pair-housed: 5 rats<br>isolated: 5 rats<br>Somatosensory isolation : 6 rats | Day 0<br>Pair-housed: $10.48 \pm 7.424$<br>Isolated.: $5.16 \pm 2.161$<br>Somatosensory isolation: $6.53 \pm 3.110$<br>Week 2<br>Pair-housed: $5 \pm 1.855$<br>Isolated.: $67.48 \pm 15.460$<br>Somatosensory isolation: $64.67 \pm 15.146$<br>Month1<br>Pair-housed: $5.08 \pm 2.488$<br>Isolated.: $67.44 \pm 23.167$<br>Somatosensory isolation: $62.4 \pm 26.582$ | RM 2_way Anova + Tukey's multiple comparisons test<br>Day 0<br>ns. $p = 0.9927$: Isolated vs. Pair-housed<br>ns. $p = 0.9999$: Isolated vs. Somatosensory isolation<br>ns. $p = 0.9966$: Pair-housed vs. Somatosensory isolation<br>Week 2<br>*$p = 0.0182$: Isolated vs. Pair-housed<br>ns. $p = 0.9987$: Isolated vs. Somatosensory isolation<br>*$p = 0.0186$: Pair-housed vs. Somatosensory isolation<br>Month1<br>*$p = 0.0185$: Isolated vs. Pair-housed<br>ns. $p = 0.9930$: Isolated vs. Somatosensory isolation<br>*$p = 0.0249$: Pair-housed vs. Somatosensory isolation |
| Fig. 1 G | 47 physical touch collected from 3 rats | Pre-social activity: $-6.124 \pm 44.83$<br>During physical touch: $586.7 \pm 179.1$ | Wilcoxon test<br>***$p = 0.0007$: Pre-social activity vs. During physical touch |
| Fig. 1 H | 28 moving away collected from 3 rats | Social proximity: $-2.504 \pm 44.78$<br>During moving away: $-388.0 \pm 114.5$ | Paired Student's T-test<br>**$p = 0.0034$: Social proximity vs. During moving away |
| Fig. 2 B1 | 8 male Wistar rat | 1st control<br>Body sniffing: $66.725 \pm 20.54$<br>Anogenital sniffing: $13.425 \pm 4.607$<br>Positive valence contact: $73.925 \pm 13.717$<br>Aggressive behavior: $150.775 \pm 32.675$<br>CNO:<br>Body sniffing: $43.575 \pm 7.67$<br>Anogenital sniffing: $6.525 \pm 2.224$ | RM 2_way Anova + Tukey's multiple comparisons test<br>Body sniffing<br>ns. $p = 0.7747$: 1st control vs. CNO<br>ns. $p = 0.9980$: 1st control vs. 2nd control<br>ns. $p = 0.7392$: CNO vs. 2nd control<br>Anogenital sniffing<br>ns. $p = 0.9775$: 1st control vs. CNO<br>ns. $p = 0.9914$: 1st control vs. 2nd control<br>ns. $p = 0.9966$: CNO vs. 2nd control<br>Positive valence contacts:<br>***$p < 0.0001$: 1st control vs. CNO<br>ns. $p > 0.9999$: 1st control vs. 2nd control |
| | | Positive valence contact:<br>$235.4 \pm 45.52$<br>Aggressive behavior:<br>$42.725 \pm 18.55$<br>2nd control:<br>Body sniffing:<br>$68.775 \pm 14.75$<br>Anogenital sniffing: $9.175 \pm 2.796$<br>Positive valence contact:<br>$74.275 \pm 18.711$<br>Aggressive behavior:<br>$159.675 \pm 49.607$ | ***p < 0.0001: 2nd control vs. CNO<br>Aggressive behavior:<br>**p = 0.0066: 1st control vs. CNO<br>ns. p = 0.9628: 1st control vs. 2nd control<br>**p = 0.0031: 2nd control vs. CNO |
| Fig. 2<br>B3 | 8 male Wistar<br>rat | 1st control<br>Dominant mounting: $65.6 \pm 16.284$<br>Fighting: $79.25 \pm 20.381$<br>Biting: $5.925 \pm 2.559$<br>CNO:<br>Dominant mounting: $26.95 \pm 15.522$<br>Fighting: $12.575 \pm 6.396$<br>Biting: $3.2 \pm 1.901$<br>2nd control:<br>Dominant mounting: $64 \pm 26.101$<br>Fighting: $92.125 \pm 30.102$<br>Biting: $3.55 \pm 1.240$ | RM 2_way Anova + Tukey's multiple<br>comparisons test<br>Dominant mounting<br>ns. p = 0.2298: 1st control vs. CNO<br>ns. p = 0.9974: 1st control vs. 2nd control<br>ns. p = 0.2578: CNO vs. 2nd control"<br>Fighting:<br>*p = 0.0170: 1st control vs. CNO<br>ns. p = 0.8444: 1st control vs. 2nd control<br>**p = 0.0038: 2nd control vs. CNO<br>Biting<br>ns. p = 0.9924: 1st control vs. CNO<br>ns. p = 0.9942: 1st control vs. 2nd control<br>ns. p = 0.9999: CNO vs. 2nd control |
| Fig. 2<br>B4 | Vehicle: 4 rats<br>CNO: 5 rats | Vehicle: $15.50 \pm 1.848$<br>CNO: $12.60 \pm 2.909$ | Unpaired Student's T-test<br>ns. p = 0.4561: vehicle vs. CNO |
| Fig. 2<br>C1 | 10 male Wistar<br>rat | 1st control<br>Body sniffing: $85.64 \pm 12.309$<br>Anogenital sniffing: $14.76 \pm 4.734$<br>Positive valence contact:<br>$102.02 \pm 7.354$<br>Aggressive behavior: $19.98 \pm 4.495$<br>CNO:<br>Body sniffing: $81.74 \pm 17.478$<br>Anogenital sniffing: $30.9 \pm 8.456$ | RM 2_way Anova + Tukey's multiple<br>comparisons test<br>Body sniffing<br>ns. p = 0.9507: 1st control vs. CNO<br>ns. p = 0.1153: 1st control vs. 2nd control<br>ns. p = 0.2069: CNO vs. 2nd control<br>Anogenital sniffing<br>ns. p = 0.4256: 1st control vs. CNO<br>ns. p = 0.3195: 1st control vs. 2nd control<br>ns. p = 0.9784: CNO vs. 2nd control<br>Positive valence contacts:<br>***p = 0.0003: 1st control vs. CNO |
|  |  | Positive valence contact:<br>48.84 ± 12.193<br>Aggressive behavior: 72.38 ± 15.915<br>2nd control:<br>Body sniffing: 59.68 ± 9.395<br>Anogenital sniffing: 33.46 ± 7.963<br>Positive valence contact:<br>99.78 ± 31.969<br>Aggressive behavior: 31.26 ± 9.752 | ns. p = 0.9834: 1st control vs. 2nd control<br>***p = 0.0005: 2nd control vs. CNO<br>Aggressive behavior:<br>***p = 0.0003: 1st control vs. CNO<br>ns. p = 0.6567: 1st control vs. 2nd control<br>**p = 0.0058: 2nd control vs. CNO |
| Fig. 2 C3 | 10 male Wistar rat | 1st control<br>Dominant mounting: 10.66 ± 2.171<br>Fighting: 8.96 ± 2.454<br>Biting: 0.36 ± 0.176<br>CNO:<br>Dominant mounting: 35.7 ± 5.028<br>Fighting: 34.68 ± 11.922<br>Biting: 2 ± 1.342<br>2nd control:<br>Dominant mounting: 16.78 ± 4.904<br>Fighting: 12.78 ± 4.432<br>Biting: 1.7 ± 0.839 | RM 2_way Anova + Tukey's multiple comparisons test<br>Dominant mounting<br>***p = 0.0007: 1st control vs. CNO<br>ns. p = 0.6046: 1st control vs. 2nd control<br>*p = 0.0121: 2nd control vs. CNO<br>Fighting:<br>***p = 0.0005: 1st control vs. CNO<br>ns. p = 0.8209: 1st control vs. 2nd control<br>**p = 0.0032: 2nd control vs. CNO<br>Biting<br>ns. p = 0.9924: 1st control vs. CNO<br>ns. p = 0.9759: 1st control vs. 2nd control<br>ns. p = 0.9999: CNO vs. 2nd control |
| Fig. 2 C4 | Vehicle: 4 rats<br>CNO: 5 rats | Vehicle: 17.25 ± 2.869<br>CNO: 14.20 ± 2.267 | Unpaired Student's T-test<br>ns. p = 0.4248: vehicle vs. CNO |
| Fig. 2 D1 | 6 male Wistar rat | 1st control<br>Body sniffing: 61.57 ± 10.222<br>Anogenital sniffing: 6.00 ± 2.836<br>Positive valence contact: 54.13 ± 10.623<br>Aggressive behavior: 25.53 ± 7.851<br>CNO:<br>Body sniffing: 72.07 ± 12.373<br>Anogenital sniffing: 23.10 ± 10.469<br>Positive valence contact: 60.30 ± 15.850 | RM 2_way Anova + Tukey's multiple comparisons test<br>Body sniffing<br>ns. p = 0.5927: 1st control vs. CNO<br>ns. p = 0.0647: 1st control vs. 2nd control<br>ns. p = 0.3837: CNO vs. 2nd control<br>Anogenital sniffing<br>ns. p = 0.2581: 1st control vs. CNO<br>ns. p = 0.1364: 1st control vs. 2nd control<br>ns. p = 0.9318: CNO vs. 2nd control<br>Positive valence contacts:<br>ns. p = 0.8334: 1st control vs. CNO<br>ns. p = 0.9741: 1st control vs. 2nd control<br>ns. p = :0.7086 2nd control vs. CNO |
| | | Aggressive behavior: $26.90 \pm 11.296$<br>2nd control:<br>Body sniffing: $86.37 \pm 16.371$<br>Anogenital sniffing: $26.93 \pm 13.483$<br>Positive valence contact: $51.80 \pm 14.369$<br>Aggressive behavior: $27.97 \pm 10.305$ | Aggressive behavior:<br>ns. $p = 0.9910$: 1st control vs. CNO<br>ns. $p = 0.9719$: 1st control vs. 2nd control<br>ns. $p = 0.9945$: 2nd control vs. CNO |
| Fig. 2 D3 | 6 male Wistar rat | 1st control<br>Dominant mounting: $10.833 \pm 6.199$<br>Fighting: $6.833 \pm 2.362$<br>Biting: $7.867 \pm 3.740$<br>CNO:<br>Dominant mounting: $13.867 \pm 9.531$<br>Fighting: $6.800 \pm 2.654$<br>Biting: $6.233 \pm 3.068$<br>2nd control:<br>Dominant mounting: $14.200 \pm 6.956$<br>Fighting: $9.833 \pm 3.295$<br>Biting: $3.933 \pm 1.903$ | RM 2_way Anova + Tukey's multiple comparisons test<br>Dominant mounting<br>ns. $p = 0.5859$: 1st control vs. CNO<br>ns. $p = 0.5192$: 1st control vs. 2nd control<br>ns. $p = 0.9934$: 2nd control vs. CNO<br>Fighting:<br>ns. $p > 0.9999$: 1st control vs. CNO<br>ns. $p = 0.5927$: 1st control vs. 2nd control<br>ns. $p = 0.5859$: CNO vs. 2nd control<br>Biting<br>ns. $p = 0.8545$: 1st control vs. CNO<br>ns. $p = 0.4119$: 1st control vs. 2nd control<br>ns. $p = 0.7335$: 2nd control vs. CNO |
| Fig. 2 D4 | Vehicle: 3 rats<br>CNO: 3 rats | Vehicle: $15.67 \pm 2.728$<br>CNO: $18.67 \pm 2.667$ | Unpaired Student's T-test<br>ns. $p = 0.4756$: vehicle vs. CNO |
| Fig. 2 E2 | Light off: 84 attacks<br>Light on: 14 attacks | Light off: $5.676 \pm 0.4998$<br>Light on: $0.2286 \pm 0.1290$ | Mann Whitney test<br>***$p < 0.0001$: light off vs. light on |
| Fig. 2 E3 | 3 male CD1 mice | Light off: $5.731 \pm 0.4767$<br>Light on: $0.2667 \pm 0.2186$ | Paired Student's T-test<br>*$p = 0.0145$: light off vs. light on |
| Fig. 2 E4 | Light off: 79 durations between attacks<br>Light on: 13 durations attacks | Light off: $13.08 \pm 2.588$<br>Light on: $26.43 \pm 8.534$ | Mann Whitney test<br>**$p = 0.0052$: light off vs. light on |
| Fig. 3<br>F | 8 male Wistar<br>rat | 1st control: $19.43 \pm 4.697$<br>CNO: $4.775 \pm 2.396$<br>2nd control: $16.85 \pm 4.312$ | Friedmann test + Dunn's multiple<br>comparisons test for aggressive behavior<br>*p = 0.0374: 1st control vs. CNO<br>*p = 0.0179: 2nd control vs. CNO |
| Fig. 3<br>H | 8 male Wistar<br>rat | 1st control<br>Dominant mounting:<br>$13.025 \pm 4.219$<br>Fighting: $5.525 \pm 0.820$<br>Biting: $0.875 \pm 0.264$<br>CNO:<br>Dominant mounting: $0.4 \pm 0.210$<br>Fighting: $1.9 \pm 0.643$<br>Biting: $2.475 \pm 2.167$<br>2nd control:<br>Dominant mounting: $6.25 \pm 2.564$<br>Fighting: $8.25 \pm 1.889$<br>Biting: $2.35 \pm 1.111$ | RM 2_way Anova + Tukey's multiple<br>comparisons test<br>Dominant mounting<br>*p = 0.0002: 1st control vs. CNO<br>ns. p = 0.0542: 1st control vs. 2nd control<br>ns. p = 0.1091: 2nd control vs. CNO<br>Fighting:<br>ns. p = 0.4137: 1st control vs. CNO<br>ns. p = 0.6041: 1st control vs. 2nd control<br>ns. p = 0.0755: CNO vs. 2nd control<br>Biting<br>ns. p = 0.8392: 1st control vs. CNO<br>ns. p = 0.8614: 1st control vs. 2nd control<br>ns. p = 0.9989: 2nd control vs. CNO |
| Fig. 3<br>I | 7 male Wistar<br>rat | 1st control<br>Body sniffing: $30.686 \pm 1.776$<br>Anogenital sniffing: $4.657 \pm 2.180$<br>Positive valence contact:<br>$45.543 \pm 9.719$<br>Aggressive behavior: $7.343 \pm 2.380$<br>CNO:<br>Body sniffing: $33.857 \pm 8.507$<br>Anogenital sniffing: $6.571 \pm 3.100$<br>Positive valence contact:<br>$12.229 \pm 6.601$<br>Aggressive behavior:<br>$26.114 \pm 7.161$<br>2nd control:<br>Body sniffing:<br>$27.400 \pm 7.049$<br>Anogenital sniffing: $3.771 \pm 2.060$<br>Positive valence contact:<br>$31.200 \pm 5.776$<br>Aggressive behavior: $4.943 \pm 1.114$ | RM 2_way Anova + Tukey's multiple<br>comparisons test<br>Body sniffing<br>ns. p = 0.9087: 1st control vs. CNO<br>ns. p = 0.9023: 1st control vs. 2nd control<br>ns. p = 0.6743: CNO vs. 2nd control<br>Anogenital sniffing<br>ns. p = 0.9657: 1st control vs. CNO<br>ns. p = 0.9925: 1st control vs. 2nd control<br>ns. p = 0.9280: CNO vs. 2nd control<br>Positive valence contacts:<br>***p = 0.0002: 1st control vs. CNO<br>ns. p = 0.1534: 1st control vs. 2nd control<br>*p = 0.0417: 2nd control vs. CNO<br>Aggressive behavior:<br>*p = 0.0444: 1st control vs. CNO<br>ns. p = 0.9466: 1st control vs. 2nd control<br>*p = 0.0205: 2nd control vs. CNO |
| Fig. 3<br>K | 7 male Wistar<br>rat | 1st control<br>Dominant mounting: $3.143 \pm 1.769$<br>Fighting: $3.743 \pm 1.231$<br>Biting: $0.457 \pm 0.253$<br>CNO:<br>Dominant mounting:<br>$12.143 \pm 6.150$<br>Fighting: $11.943 \pm 4.758$<br>Biting: $2.029 \pm 1.012$<br>2nd control:<br>Dominant mounting: $1.857 \pm 0.759$<br>Fighting: $2.686 \pm 0.881$<br>Biting: $0.400 \pm 0.175$ | RM 2_way Anova + Tukey's multiple<br>comparisons test<br>Dominant mounting<br>*p = 0.0438: 1st control vs. CNO<br>ns. p = 0.9321: 1st control vs. 2nd control<br>*p = 0.0187: 2nd control vs. CNO<br>Fighting:<br>ns. p = 0.0716: 1st control vs. CNO<br>ns. p = 0.9535: 1st control vs. 2nd control<br>*p = 0.0372: 2nd control vs. CNO<br>Biting<br>ns. p = 0.9004: 1st control vs. CNO<br>ns. p = 0.9999: 1st control vs. 2nd control<br>ns. p = 0.8935: 2nd control vs. CNO |
| Fig. 4<br>A | Vehicle: 5 rats<br>CNO: 4 rats | Vehicle:<br>PIL: $111.5 \pm 8.104$<br>MPOA: $279.6 \pm 33.361$<br>PVN: $304.4 \pm 45.087$<br>DMH: $244.6 \pm 23.105$<br>MeA: $143.3 \pm 15.499$<br>PAG: $186.9 \pm 34.696$<br>CNO:<br>PIL: $292.125 \pm 37.406$<br>MPOA: $502.125 \pm 65.689$<br>PVN: $303.5 \pm 117.628$<br>DMH: $315.5 \pm 43.394$<br>MeA: $196.375 \pm 14.595$<br>PAG: $174.875 \pm 41.261$ | 2_way Anova + Sidak's multiple<br>comparisons test<br>PIL:<br>*p = 0.0434: vehicle vs. CNO<br>MPOA:<br>**p = 0.0073: vehicle vs. CNO<br>PVN:<br>ns. p > 0.9999: vehicle vs. CNO<br>DMH:<br>ns. p = 0.8551: vehicle vs. CNO<br>MeA:<br>ns. p = 0.9589: vehicle vs. CNO<br>PAG<br>ns. p > 0.9999: vehicle vs. CNO |
| Fig. 4<br>E | Vehicle: 4 rats<br>CNO: 4 rats | Vehicle:<br>PIL: $213.875 \pm 20.526$<br>MPOA: $616.5 \pm 43.521$<br>PVN: $368.875 \pm 57.421$<br>DMH: $284.75 \pm 19.335$<br>MeA: $215.625 \pm 32.669$<br>PAG: $158.875 \pm 29.619$<br>CNO:<br>PIL: $91.5 \pm 4.113$<br>MPOA: $250.75 \pm 10.217$<br>PVN: $204.5 \pm 16.596$ | 2_way Anova + Sidak's multiple<br>comparisons test<br>PIL:<br>*p = 0.0248: vehicle vs. CNO<br>MPOA:<br>***p < 0.0001: vehicle vs. CNO<br>PVN:<br>**p < 0.0013: vehicle vs. CNO<br>DMH:<br>ns. p = 0.3989: vehicle vs. CNO |
| | | DMH: $213 \pm 22.546$<br>MeA: $198.25 \pm 22.629$<br>PAG: $92.5 \pm 16.023$ | MeA:<br>ns. $p = 0.9986$: vehicle vs. CNO<br>PAG<br>ns. $p = 0.4888$: vehicle vs. CNO |
| Fig. 4 C | 10 male Wistar rat | 1st control<br>Body sniffing: $82.520 \pm 9.882$<br>Anogenital sniffing: $23.920 \pm 7.138$<br>Positive valence contact: $96.640 \pm 21.878$<br>Aggressive behavior: $73.920 \pm 14.416$<br>CNO:<br>Body sniffing: $87.260 \pm 10.901$<br>Anogenital sniffing: $23.560 \pm 4.220$<br>Positive valence contact: $243.480 \pm 41.039$<br>Aggressive behavior: $14.000 \pm 3.302$<br>2nd control:<br>Body sniffing: $76.020 \pm 11.388$<br>Anogenital sniffing: $32.860 \pm 10.442$<br>Positive valence contact: $96.480 \pm 26.872$<br>Aggressive behavior: $56.560 \pm 16.713$ | RM 2_way Anova + Tukey's multiple comparisons test<br>Body sniffing<br>ns. $p = 0.9751$: 1st control vs. CNO<br>ns. $p = 0.9537$: 1st control vs. 2nd control<br>ns. $p = 0.8679$: CNO vs. 2nd control<br>Anogenital sniffing<br>ns. $p = 0.9999$: 1st control vs. CNO<br>ns. $p = 0.9142$: 1st control vs. 2nd control<br>ns. $p = 0.9075$: CNO vs. 2nd control<br>Positive valence contacts:<br>***$p < 0.0001$: 1st control vs. CNO<br>ns. $p > 0.9999 = 0.9075$: 1st control vs. 2nd control<br>***$p < 0.0001$: 2nd control vs. CNO<br>Aggressive behavior:<br>*$p = 0.0229$: 1st control vs. CNO<br>ns. $p = 0.7141$: 1st control vs. 2nd control<br>ns. $p = 0.1399$: 2nd control vs. CNO |
| Fig. 4 D | 4 male Wistar rat | 1st control<br>Body sniffing: $71.100 \pm 7.465$<br>Anogenital sniffing: $69.400 \pm 12.789$<br>Positive valence contact: $128.450 \pm 22.846$<br>Aggressive behavior: $37.050 \pm 4.073$<br>CNO:<br>Body sniffing: $60.250 \pm 3.640$<br>Anogenital sniffing: $77.850 \pm 9.624$<br>Positive valence contact: $55.500 \pm 8.980$ | RM 2_way Anova + Tukey's multiple comparisons test<br>Body sniffing<br>ns. $p = 0.8753$: 1st control vs. CNO<br>ns. $p = 0.6653$: 1st control vs. 2nd control<br>ns. $p = 0.9257$: CNO vs. 2nd control<br>Anogenital sniffing<br>ns. $p = 0.9222$: 1st control vs. CNO<br>ns. $p = 0.6499$: 1st control vs. 2nd control<br>ns. $p = 0.8677$: CNO vs. 2nd control<br>Positive valence contacts:<br>**$p = 0.0079$: 1st control vs. CNO<br>ns. $p = 0.9982$: 1st control vs. 2nd control<br>**$p = 0.0090$: 2nd control vs. CNO |
|  |  | Aggressive behavior:<br>127.600 ± 20.467<br>2nd control:<br>Body sniffing: 52.000 ± 6.761<br>Anogenital sniffing: 89.050 ± 19.815<br>Positive valence contact:<br>127.200 ± 20.758<br>Aggressive behavior:<br>37.000 ± 12.296 | Aggressive behavior:<br>**p = 0.0011: 1st control vs. CNO<br>ns. p > 0.9999: 1st control vs. 2nd control<br>**p = 0.0011: 2nd control vs. CNO |
| Fig. 5 F | 7 male Wistar rat | 1st control<br>Body sniffing: 31.629 ± 3.760<br>Anogenital sniffing: 70.743 ± 13.996<br>Positive valence contact:<br>108.057 ± 26.403<br>Aggressive behavior:<br>15.343 ± 3.697<br>CNO:<br>Body sniffing: 22.886 ± 3.820<br>Anogenital sniffing: 68.086 ± 17.101<br>Positive valence contact:<br>31.000 ± 7.585<br>Aggressive behavior:<br>70.600 ± 13.360<br>2nd control:<br>Body sniffing:<br>25.629 ± 2.701<br>Anogenital sniffing: 69.743 ± 19.424<br>Positive valence contact:<br>80.743 ± 18.463<br>Aggressive behavior:<br>19.800 ± 4.019 | RM 2_way Anova + Tukey's multiple comparisons test<br>Body sniffing<br>ns. p = 0.8433: 1st control vs. CNO<br>ns. p = 0.9227: 1st control vs. 2nd control<br>ns. p = 0.9833: CNO vs. 2nd control<br>Anogenital sniffing<br>ns. p = 0.9843: 1st control vs. CNO<br>ns. p = 0.9978: 1st control vs. 2nd control<br>ns. p = 0.9939: CNO vs. 2nd control<br>Positive valence contacts:<br>***p < 0.0001: 1st control vs. CNO<br>ns. p = 0.2008: 1st control vs. 2nd control<br>**p = 0.0074: 2nd control vs. CNO<br>Aggressive behavior:<br>**p = 0.0027: 1st control vs. CNO<br>ns. p = 0.9565: 1st control vs. 2nd control<br>**p = 0.0061: 2nd control vs. CNO |
| Fig. 5 H | 7 male Wistar rat | 1st control<br>Dominant mounting: 7.829 ± 2.603<br>Fighting: 3.429 ± 0.539<br>Biting: 4.086 ± 1.388<br>CNO:<br>Dominant mounting:<br>33.771 ± 9.296<br>Fighting: 29.057 ± 7.844<br>Biting: 7.771 ± 2.359 | RM 2_way Anova + Tukey's multiple comparisons test<br>Dominant mounting<br>***p < 0.0001: 1st control vs. CNO<br>ns. p = 0.3584: 1st control vs. 2nd control<br>**p = 0.0023: 2nd control vs. CNO<br>Fighting:<br>***p < 0.0001: 1st control vs. CNO<br>ns. p = 0.9849: 1st control vs. 2nd control |
| | | 2nd control:<br>Dominant mounting:<br>$14.971 \pm 3.584$<br>Fighting: $4.286 \pm 1.073$<br>Biting: $0.543 \pm 0.275$ | ***p < 0.0001: 2nd control vs. CNO<br>Biting<br>ns. p = 0.7560: 1st control vs. CNO<br>ns. p = 0.7721: 1st control vs. 2nd control<br>ns. p = 0.3499: 2nd control vs. CNO |
| Fig. 5 J | Vehicle: 4 rats<br>CNO: 4 rats | Vehicle:<br>PIL: $499.875 \pm 55.106$<br>VMH: $406.75 \pm 61.412$<br>CeA: $449.25 \pm 31.768$<br>PVT: $516.75 \pm 19.712$<br>PVN: $309 \pm 58.793$<br>MPOA: $706.25 \pm 75.991$<br>LS: $385 \pm 21.543$<br>CNO:<br>PIL: $255 \pm 49.434$<br>VMH: $860.5 \pm 70.365$<br>CeA: $405.125 \pm 22.512$<br>PVT: $485.5 \pm 40.975$<br>PVN: $366.625 \pm 55.275$<br>MPOA: $806.5 \pm 22.576$<br>LS: $475.25 \pm 16.468$ | 2_way Anova + Sidak's multiple comparisons test<br>PIL:<br>*p = 0.0049: vehicle vs. CNO<br>VMH:<br>***p < 0.0001: vehicle vs. CNO<br>CeA:<br>ns. p = 0.9935: vehicle vs. CNO<br>PVT:<br>ns. p = 0.9993: vehicle vs. CNO<br>PVN:<br>ns. p = 0.9699: vehicle vs. CNO<br>MPOA:<br>ns. p = 0.6561: vehicle vs. CNO<br>LS:<br>ns. p = 0.7601: vehicle vs. CNO |
| Fig. S2 A | Vehicle: 4 rats<br>CNO: 5 rats | Vehicle: $239.7 \pm 41.55$<br>CNO: $247.3 \pm 26.98$ | Unpaired Student's T-test<br>ns. p = 0.8770: vehicle vs. CNO |
| Fig. S2 B | Vehicle: 4 rats<br>CNO: 5 rats | Vehicle: $193.0 \pm 34.25$<br>CNO: $184.6 \pm 37.46$ | Unpaired Student's T-test<br>ns. p = 0.8770: vehicle vs. CNO |
| Fig. S2 C | Vehicle: 3 rats<br>CNO: 3 rats | Vehicle: $244.5 \pm 58.36$<br>CNO: $217.3 \pm 30.04$ | Unpaired Student's T-test<br>ns. p = 0.6992: vehicle vs. CNO |
| Fig. S2 D | Vehicle: 6 rats<br>CNO: 5 rats | Vehicle: $84.23 \pm 36.50$<br>CNO: $159.0 \pm 33.90$ | Mann Whitney test<br>ns. p = 0.0519: vehicle vs. CNO |
| Fig. S2 E | Vehicle: 4 rats<br>CNO: 4 rats | Vehicle: $205.3 \pm 58.72$<br>CNO: $94.90 \pm 8.928$ | Unpaired Student's T-test<br>ns. p = 0.1124: vehicle vs. CNO |
| Fig. S2 E | Vehicle: 3 rats<br>CNO: 4 rats | Vehicle: $276.8 \pm 111.9$<br>CNO: $424.1 \pm 51.89$ | Unpaired Student's T-test<br>ns. p = 0.2455: vehicle vs. CNO |
| Fig. S2 F | Vehicle: 4 rats<br>CNO: 4 rats | Vehicle: $337.1 \pm 28.74$<br>CNO: $281.0 \pm 67.50$ | Unpaired Student's T-test<br>ns. p = 0.4735: vehicle vs. CNO |
| Fig. S2<br>G | Vehicle: 6 rats<br>CNO: 5 rats | Vehicle: $9.667 \pm 2.753$<br>CNO: $10.80 \pm 2.538$ | Unpaired Student's T-test<br>ns. $p = 0.7728$: vehicle vs. CNO |
| Fig. S2<br>H | Vehicle: 4 rats<br>CNO: 4 rats | Vehicle: $14.25 \pm 3.119$<br>CNO: $8.500 \pm 1.443$ | Unpaired Student's T-test<br>ns. $p = 0.1454$: vehicle vs. CNO |
| Fig. S2<br>I | Vehicle: 3 rats<br>CNO: 4 rats | Vehicle: $16.33 \pm 2.955$<br>CNO: $29.00 \pm 3.342$ | Unpaired Student's T-test<br>*$p = 0.0421$: vehicle vs. CNO |
| Fig. S2<br>H | Vehicle: 4 rats<br>CNO: 4 rats | Vehicle: $15.75 \pm 28.74$<br>CNO: $12.50 \pm 2.754$ | Unpaired Student's T-test<br>ns. $p = 0.4517$: vehicle vs. CNO |
| Fig. S4<br>A | 10 male Wistar<br>rat | 1st control<br>Dominant mounting: $37.78 \pm 8.37$<br>Fighting: $19.48 \pm 3.61$<br>Biting: $16.66 \pm 6.20$<br>CNO:<br>Dominant mounting: $7.62 \pm 2.48$<br>Fighting: $3.2 \pm 0.71$<br>Biting: $3.18 \pm 0.85$<br>2nd control:<br>Dominant mounting: $34.1 \pm 12.54$<br>Fighting: $16.66 \pm 2.04$<br>Biting: $5.8 \pm 1.52$ | RM 2_way Anova + Tukey's multiple<br>comparisons test<br>Dominant mounting<br>***$p = 0.0003$: 1st control vs. CNO<br>ns. $p = 0.8631$: 1st control vs. 2nd control<br>**$p = 0.0013$: 2nd control vs. CNO<br>Fighting:<br>ns. $p = 0.0659$: 1st control vs. CNO<br>ns. $p = 0.9171$: 1st control vs. 2nd control<br>ns. $p = 0.1505$: 2nd control vs. CNO<br>Biting<br>ns. $p = 0.1497$: 1st control vs. CNO<br>ns. $p = 0.2864$: 1st control vs. 2nd control<br>ns. $p = 0.9280$: 2nd control vs. CNO |
| Fig. S9<br>C | 4 male Wistar<br>rat | 1st control<br>Dominant mounting: $29.95 \pm 6.83$<br>Fighting: $5.55 \pm 2.43$<br>Biting: $1.55 \pm 1.16$<br>CNO:<br>Dominant mounting: $103.35 \pm 27.96$<br>Fighting: $19.35 \pm 2.39$<br>Biting: $4.9 \pm 2.03$<br>2nd control:<br>Dominant mounting: $30.65 \pm 15.41$<br>Fighting: $5.6 \pm 1.56$<br>Biting: $0.75 \pm 0.75$ | RM 2_way Anova + Tukey's multiple<br>comparisons test<br>Dominant mounting<br>***$p < 0.0001$: 1st control vs. CNO<br>ns. $p = 0.9979$: 1st control vs. 2nd control<br>***$p < 0.0001$: 2nd control vs. CNO<br>Fighting:<br>ns. $p = 0.4641$: 1st control vs. CNO<br>ns. $p > 0.9999$: 1st control vs. 2nd control<br>ns. $p = 0.4665$: 2nd control vs. CNO<br>Biting<br>ns. $p = 0.9539$: 1st control vs. CNO<br>ns. $p = 0.9973$: 1st control vs. 2nd control<br>ns. $p = 0.9302$: 2nd control vs. CNO |
| Fig. S5<br>B | Male intruder:<br>9 rats | Male intruder:<br>PAG: $316.1 \pm 40.9$ | 2_way Anova + Sidak's multiple<br>comparisons test |
| | Control: 10 rats | PIL: $633.2 \pm 91.5$<br>DMH: $531 \pm 64.1$<br>MeA: $600.8 \pm 75.5$<br>PVN: $202.2 \pm 25.1$<br>MPOA: $595.7 \pm 89.4$<br>LS: $312.2 \pm 47.2$<br>ILC: $1175.4 \pm 164.0$<br>Control:<br>PAG: $37.1 \pm 7.3$<br>PIL: $64.4 \pm 10.1$<br>DMH: $74.4 \pm 16.7$<br>MeA: $69.8 \pm 14.0$<br>PVN: $43.7 \pm 8.4$<br>MPOA: $125.1 \pm 23.4$<br>LS: $65.5 \pm 25.4$<br>ILC: $223.5 \pm 39.7$ | PAG<br>**p = 0.0082: male intruder vs. control<br>PIL<br>***p < 0.0001: male intruder vs. control<br>DMH<br>***p < 0.0001: male intruder vs. control<br>MeA<br>***p < 0.0001: male intruder vs. control<br>PVN<br>ns. p = 0.3844: male intruder vs. control<br>MPOA<br>***p < 0.0001: male intruder vs. control<br>LS<br>*p = 0.0282: male intruder vs. control<br>ILC<br>***p < 0.0001: male intruder vs. control |

**Table S7.**
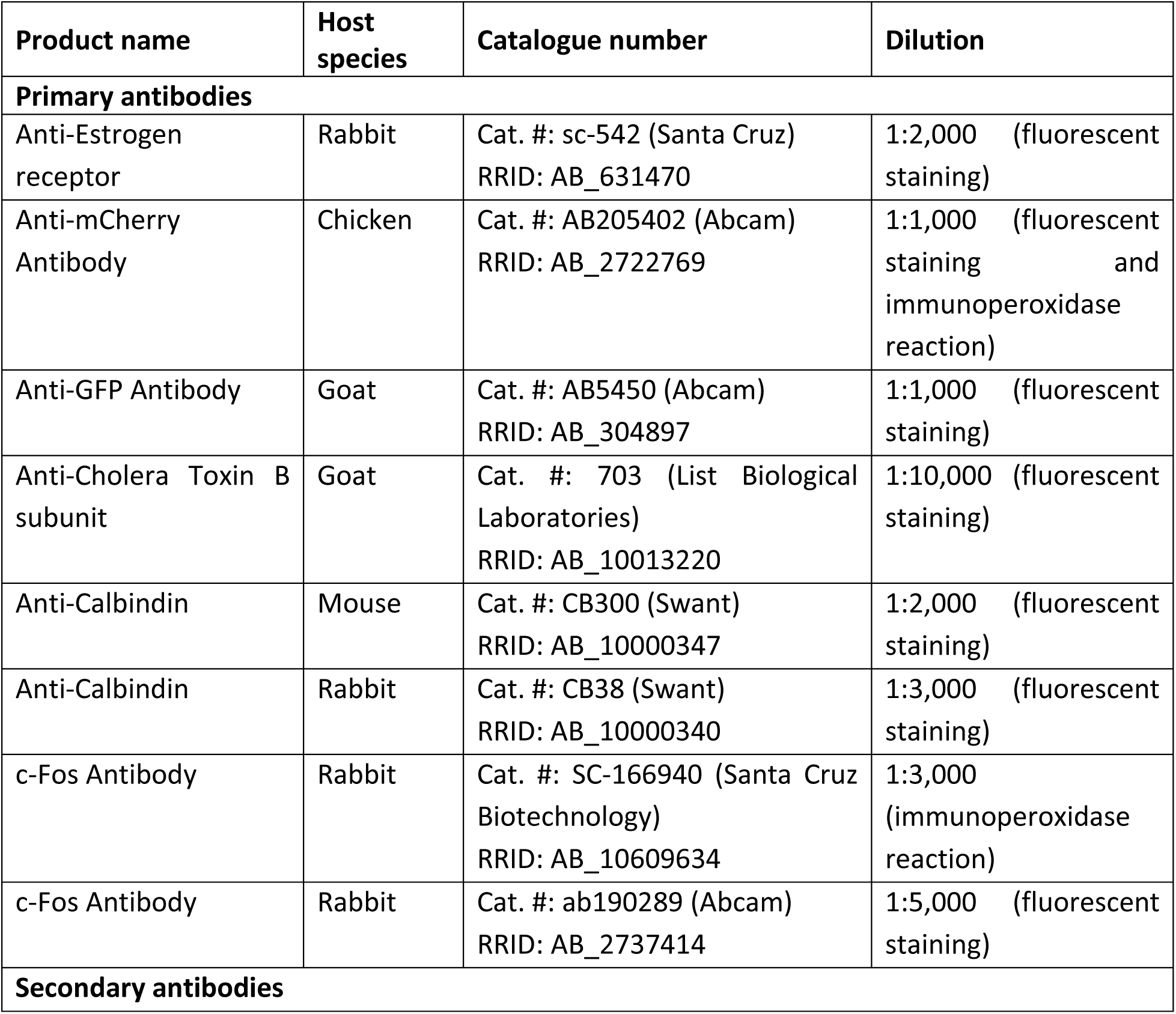
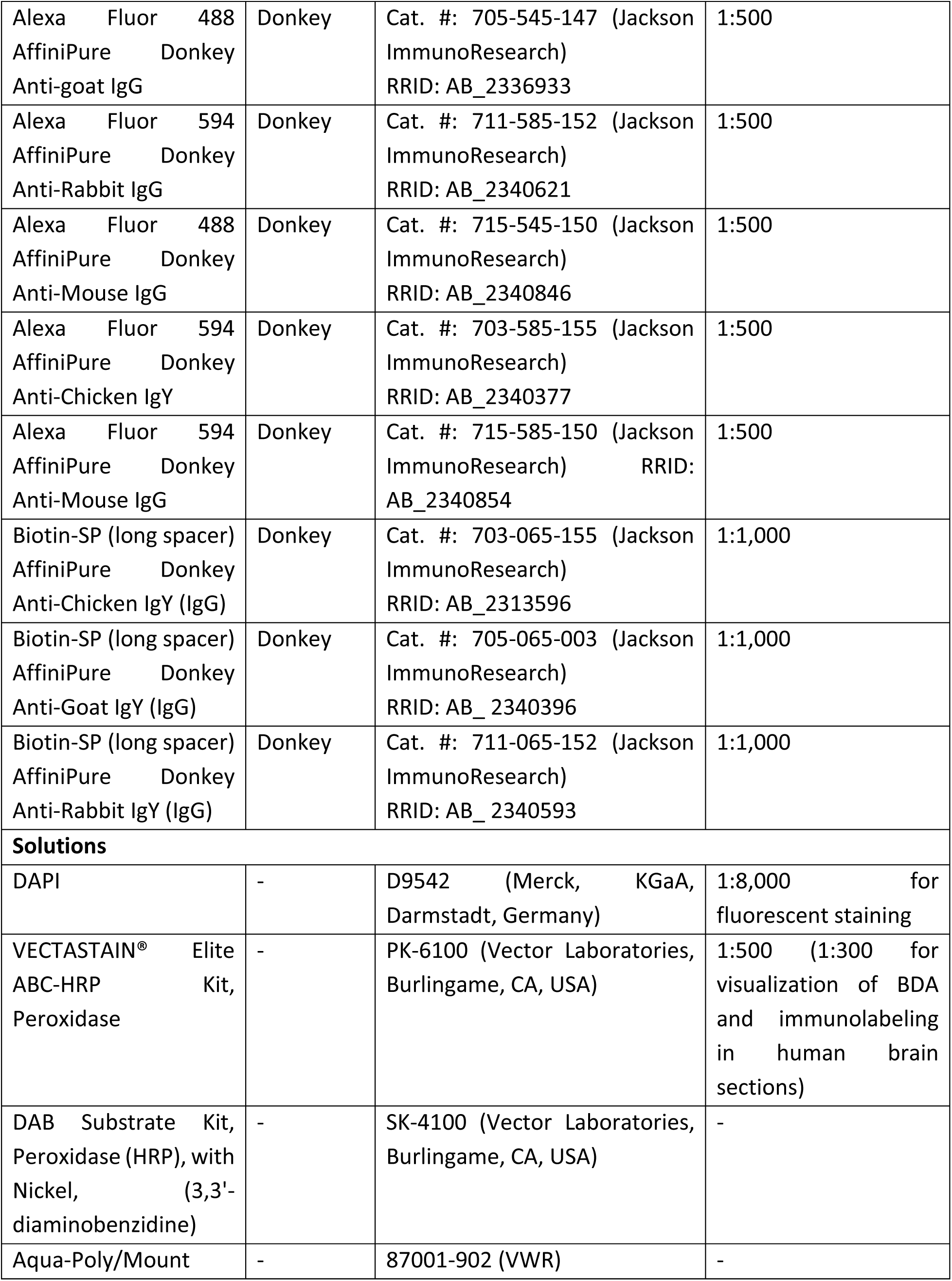
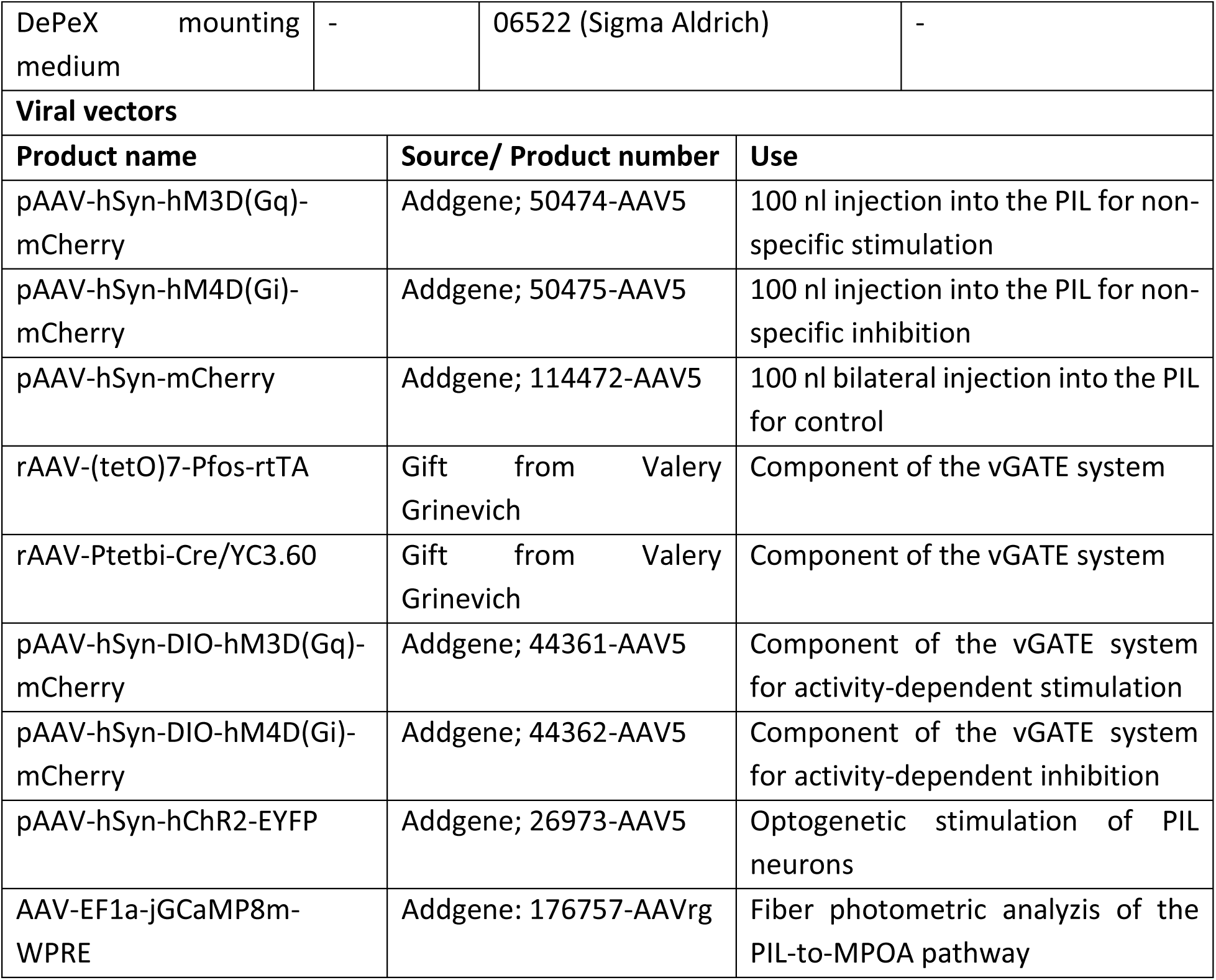
Antibodies and histological reagents used in the study.

